# A microbiota-driven neuroepithelial feed-forward circuit drives gastroesophageal junction tumorigenesis

**DOI:** 10.64898/2026.10.13.759004

**Authors:** Yi Zeng, Zaisheng Ye, Feijing Wu, Puran Zhang, Hiroki Kobayashi, Junya Arai, Xiaofei Zhi, Martin Borgmann, Jin Qian, Shuang Li, Biyun Zheng, Hualong Zheng, Ruhong Tu, Juli Lin, Masahiro Hata, Yosuke Ochiai, Quin T. Waterbury, Nagendra Harry, Leah B. Zamechek, Kara L. Vasilew, Michael Quante, Julian A. Abrams, James G. Fox, Timothy C. Wang

**Author notes:** Correspondence to: Timothy C. Wang. These authors contributed equally to this work (Yi Zeng, Zaisheng Ye, Feijing Wu, Puran Zhang).

## Abstract

The gastroesophageal junction (GEJ) is continuously exposed to refluxed bile acids and microbial perturbations, yet how these environmental cues are converted into persistent oncogenic signaling is unknown. Here we show that bile acid remodeling enriches the oral commensal *Prevotella melaninogenica* at the GEJ, where bacterial succinate activates both vagal sensory neurons and SUCNR1-expressing tuft cells. Activated tuft cells release 15-deoxy-Δ12,14-prostaglandin J2(15-d-PGJ2), which sensitizes sensory neurons and sustains nerve remodeling, while neuronal CGRP drives expansion and tuft-cell differentiation of neuro-responsive epithelial progenitors through RAMP1. This self-reinforcing microbiota–neuron– epithelium circuit promotes dysplastic progression, and its disruption—genetic or pharmacological, including with the approved CGRP antagonist rimegepant—suppresses tumor development, revealing an interceptable mechanism linking environmental dysbiosis to cancer initiation through neural circuits.

**One-sentence summary:** Bile acids enrich an oral bacterium that initiates a nerve–epithelium circuit driving gastroesophageal junction cancer.

## INTRODUCTION

The gastroesophageal junction (GEJ), where the esophageal and gastric environments converge, is an ecological transition zone continuously exposed to refluxed gastric acid and bile, resident microbiota, dietary metabolites, and chronic inflammatory stimuli (1–3). Gastroesophageal reflux disease (GERD), which affects roughly one-fifth of Western populations, chronically exposes the distal esophagus and GEJ to acid and bile and constitutes the principal risk factor for Barrett’s esophagus and esophageal adenocarcinoma, together with central obesity, male sex, and smoking (2–3). Driven by these exposures, the incidence of GEJ adenocarcinoma has risen rapidly over the past four decades, and affected patients continue to face poor outcomes despite advances in surveillance, surgical resection, and systemic therapy (1, 3). Although bile acid reflux and microbial dysbiosis have both been implicated in GEJ carcinogenesis, how these environmental cues are sensed and converted into persistent oncogenic programs within the GEJ niche remains largely unknown.

Microbial ecosystems can regulate tumor initiation not only through inflammation and immune modulation but also through metabolic remodeling of the local tissue microenvironment. In the colorectum, for example, *Fusobacterium nucleatum* is enriched in tumors and persists in metastases, and the bacterial genotoxin colibactin directly alkylates epithelial DNA (4–6). At the GEJ, the microbiota differs fundamentally from the *Helicobacter pylori*–dominated ecology of the distal stomach: profiling of the esophageal and junctional microbiome has identified distinct *Streptococcus*- and *Prevotella*-dominated signatures that track with Barrett’s esophagus and its progression to adenocarcinoma (7–9). Epidemiological studies further link periodontal disease— a hallmark of oral microbial dysbiosis—to significantly increased risks of esophageal and gastric adenocarcinoma, implicating oral microbial communities in upper gastrointestinal tumorigenesis (10–11). Bile acids, meanwhile, are established environmental drivers of Barrett’s metaplasia and can directly damage epithelial DNA (12–13). Whether bile reflux also promotes carcinogenesis indirectly—by reshaping which microbes colonize the GEJ—and how microbial and metabolic signals are then integrated into sustained oncogenic signaling remain poorly understood.

Barrier tissues monitor their chemical environment through complementary epithelial and neural sensors. Tuft cells detect luminal metabolites through specialized chemosensory receptors— including SUCNR1, which senses succinate—and secrete bioactive mediators that regulate epithelial homeostasis, immunity, and repair (14–17). Sensory neurons, in turn, directly detect bacterial products and tissue-derived danger signals, coordinating host defense, pain, and regeneration (18–19). In the intestine, enteroendocrine cells form putative synapses with vagal sensory neurons, establishing rapid epithelium-to-nerve sensory transduction (20–21), and bidirectional tuft cell–neuron communication has been proposed at mucosal barriers (17). Neural signaling is now recognized as a driver of solid tumors (22–23): denervation suppresses gastric tumorigenesis (24), nerve growth factor–driven cholinergic signaling promotes gastric cancer whereas cholinergic input restrains pancreatic cancer (25–26), glutamatergic synapses fuel glioma (27–28), and nociceptor-derived CGRP modulates cancer immunosurveillance (29). In gastric cancer specifically, nociceptive neurons engage tumor cells through a CGRP–RAMP1 axis to accelerate progression (30). Whether microbial ecosystems exploit neuroepithelial communication to initiate tumorigenesis, which epithelial populations respond to neuronal signals, and what activates these circuits in the first place remain unknown.

Here, we define a microbiota-driven neuroepithelial feed-forward circuit that links environmental dysbiosis to GEJ tumorigenesis. In human esophageal and GEJ adenocarcinoma, *TP53* mutation is the most frequent and among the earliest genetic alterations, typically arising at the transition from metaplasia to dysplasia and biasing junctional progenitors toward dysplasia rather than metaplasia (31–32). Using the *L2-IL1B* mouse model of Barrett’s-like metaplasia—in which bile acids and inflammation activate CCK2R-expressing cardia stem/progenitor cells at the squamocolumnar junction, and in which p53 mutation or high-fat diet (HFD) feeding accelerates dysplastic progression (33–35)—together with spatial imaging, single-cell transcriptomics, lineage tracing, human multi-omic profiling, and microbial colonization, we show that bile acid remodeling establishes a *Prevotella melaninogenica* niche whose succinate output activates both vagal sensory neurons and SUCNR1-expressing tuft cells. Activated tuft cells release 15-deoxy-Δ12,14-prostaglandin J2 (15-d-PGJ2), which sensitizes sensory neurons and sustains nerve remodeling, while neuronal CGRP drives expansion and tuft-cell differentiation of neuro-responsive RAMP1-high CCK2R+ progenitors through RAMP1, establishing a self-reinforcing circuit that can be disrupted genetically or pharmacologically.

## RESULTS

### Progressive expansion of tuft cell–sensory neuron niches during GEJ tumorigenesis

To characterize the cellular and spatial architecture of GEJ tumorigenesis, we performed multiplexed imaging mass cytometry on lesions from the *L2-IL1B* model, in which Barrett’s-like metaplasia at the GEJ progresses to dysplasia, is accelerated by HFD feeding (33), and preferentially accumulates tuft cells (36–37). Cell segmentation and phenotypic annotation revealed extensive remodeling of epithelial, immune, stromal, and neural compartments in HFD-accelerated lesions (*Fig. 1, A and B, and fig. S1A*). A directional neighbor-preference analysis showed a progression-associated trend toward increased tuft cell–neuron proximity, although this association did not remain significant after multiple-testing correction (*fig. S1B*). Because thin axonal and neurite processes are poorly captured by cell segmentation, we additionally analyzed β3-tubulin as a continuous image field (*fig. S1C*). Across mice, tuft-cell density positively correlated with the β3-tubulin bright-signal area fraction (Spearman ρ = 0.45, *P* = 0.012; *Fig. 1C*). Tuft cells were also significantly closer to stromal β3-tubulin–bright structures than matched non-tuft epithelial cells sampled from the same regions (*P* = 0.012; *Fig. 1C*). Compartment-resolved analysis further showed redistribution of β3-tubulin–bright signals across epithelial and stromal regions during dysplastic progression (*fig. S1D*), while area-normalized tuft-cell counts within 10, 25, and 50 μm of β3-tubulin–bright structures increased with dysplasia severity (*fig. S1E*). Consistent with these findings, the absolute number of tuft cells within 20 μm of β3-tubulin–bright pixels showed an increasing trend across dysplasia groups, although the score 1 versus score 3 comparison did not reach statistical significance (*P* = 0.130; *Fig. 1C*). Together, these analyses reveal progressive expansion of a tuft cell–neural niche during GEJ dysplastic progression.

**Fig. 1.**
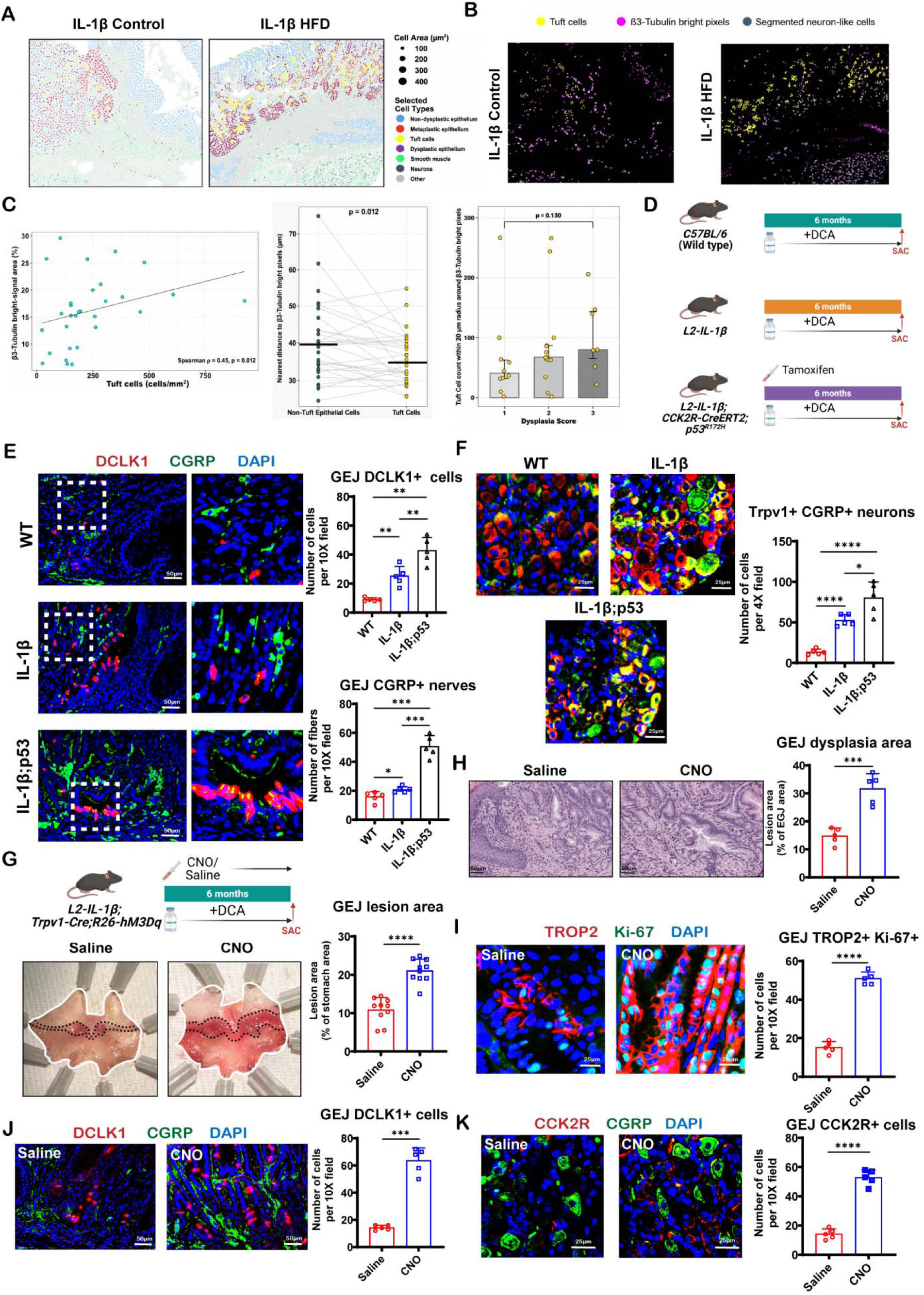
Progressive expansion of tuft cell–sensory neuron niches during GEJ tumorigenesis. (**A**) Imaging mass cytometry (IMC) cell-type maps of gastroesophageal junction (GEJ) lesions from *L2-IL1B* mice feed a control diet or high-fat diet (HFD), with segmented cells colored by phenotypic annotation. (**B**) Spatial distribution of DCLK1+ tuft cells (yellow), β3-tubulin–bright pixels (magenta), and segmented neuron-like cells (blue) in representative control- and HFD-fed *L2-IL1B* GEJ lesions. (**C**) Spatial association between tuft cells and β3-tubulin–bright neural structures. Left, correlation between tuft-cell density and β3-tubulin–bright-signal area fraction across mice (Spearman ρ = 0.45, *P* = 0.012). Middle, paired nearest distance of tuft cells and matched non-tuft epithelial cells from the same regions to stromal β3-tubulin–bright structures (*P* = 0.012). Right, number of tuft cells within 20 μm of β3-tubulin–bright pixels across dysplasia score groups; score 1 versus score 3, *P* = 0.130. (**D**) Experimental scheme: wild-type (WT), *L2-IL1B*(IL-1β), and *L2-IL1B; Cck2r-CreERT2; p53^R172H^*(IL-1β; p53) mice treated with deoxycholic acid (DCA) in drinking water for 6 months after tamoxifen induction. (**E**) Immunofluorescence (IF) of DCLK1, CGRP, and DAPI at the GEJ of WT, IL-1β, and IL-1β; p53 mice, with quantification of GEJ DCLK1+ tuft cells and CGRP+ sensory nerves (cells per 10× field). n = 5 mice per group. (**F**) IF of TRPV1 and CGRP in NG of the same genotypes, with quantification of TRPV1+CGRP+ neurons (per 4× field). n = 5 mice per group. (**G**) Scheme of chemogenetic sensory-neuron activation: *L2-IL1B; Trpv1-Cre; R26-hM3Dq* mice received clozapine-N-oxide (CNO) or saline, with DCA, for 6 months; representative whole-mount stomachs and quantification of GEJ lesion area (percentage of stomach area). n = 10 mice per group. (**H**) H&E-stained GEJ sections and quantification of dysplasia area. n = 5 mice per group. (**I**) IF of TROP2 and Ki-67 with quantification of TROP2+Ki-67+ cells. n = 5 mice per group. (**J**) IF of DCLK1 and CGRP with quantification of GEJ DCLK1+ cells. n = 5 mice per group. (**K**) IF of CCK2R and CGRP with quantification of GEJ CCK2R+ cells. n = 5 mice per group. Scale bars: 25 μm (**E, H, J**), 50 μm (**F, I, K**). For all panels, values indicate mean ± SEM. \**P* < 0.05, \*\**P* < 0.01, \*\*\**P* < 0.001, \*\*\*\**P* < 0.0001; ns: not significant. Schematic in (**D, G**) created with BioRender.com.

Vagal sensory neurons provide the predominant sensory innervation of the murine stomach and GEJ region (24–25). We therefore examined sensory innervation across wild-type, *L2-IL1B* (IL-1β), and *L2-IL1B; Cck2r-CreERT2; p53^R172H^* (IL-1β; p53) mice, which model progressively advanced disease *(Fig. 1D)*. In all groups, deoxycholic acid (DCA) was administered in drinking water to model bile reflux, as previously established for the *L2-IL1B* model (31,33). DCLK1+ tuft cells and CGRP+ sensory fibers increased stepwise from wild-type to *L2-IL1B* to p53-mutant mice *(Fig. 1E)*, and TRPV1+CGRP+ sensory neurons in the vagal nodose ganglia (NG) expanded in parallel *(Fig. 1F)*, indicating that expansion of vagal sensory innervation accompanies progressive remodeling of the tuft-cell niche.

To test whether increased sensory neuronal activity is sufficient to accelerate tumorigenesis, we chemogenetically activated TRPV1-lineage sensory neurons in *L2-IL1B; Trpv1-Cre; R26-hM3Dq* mice through chronic clozapine-N-oxide administration *(Fig. 1G)*. Compared with saline-treated controls, neuronal activation substantially increased gross lesion area and histologic dysplasia *(Fig. 1, G and H)*, expanded dysplastic TROP2+Ki-67+ epithelial cells *(Fig. 1I)*, increased DCLK1+ tuft cells together with CGRP+ innervation *(Fig. 1J)*, and enriched CCK2R+ epithelial progenitors closely apposed to CGRP+ nerve fibers *(Fig. 1K)*. Sustained activation of TRPV1-lineage sensory neurons is thus sufficient to remodel the GEJ epithelial niche and promote dysplastic progression.

### A neuro-responsive RAMP1-high CCK2R+ progenitor state mediates CGRP-dependent epithelial remodeling during GEJ tumorigenesis

To identify the epithelial source of tuft-cell expansion, we reanalyzed our previously published single-cell RNA sequencing (scRNA-seq) dataset (GSE246280) of epithelial cells from *L2-IL1B* and *L2-IL1B; Cck2r-CreERT2; p53^R172H^* mice *(fig. S2, A and B)*. Comparative clustering revealed extensive epithelial remodeling during progression: a mature tuft-cell population (cluster 9; *Dclk1, Pou2f3, Trpm5, Gnat3*) expanded in advanced lesions, and a previously unrecognized RAMP1-high (RAMP1, a key component of the CGRP receptor) CCK2R+ population (cluster 11) emerged and expanded in parallel *(Fig. 2, A and B, and fig. S2, C to E)*. This progenitor state lay immediately adjacent to mature tuft cells in transcriptional space (Fig. 2C), and pseudotime analysis placed it upstream of tuft cells, with progressive acquisition of the tuft-cell transcriptional program along the differentiation trajectory *(Fig. 2D and fig. S2, F and G)*. Tuft cells from advanced lesions exhibited enhanced expression of canonical effector genes, including *Hpgds*, *Ptgs1*, and *Il25 (Fig. 2E)*, indicating acquisition of a functionally mature phenotype during tumor progression.

**Fig. 2.**
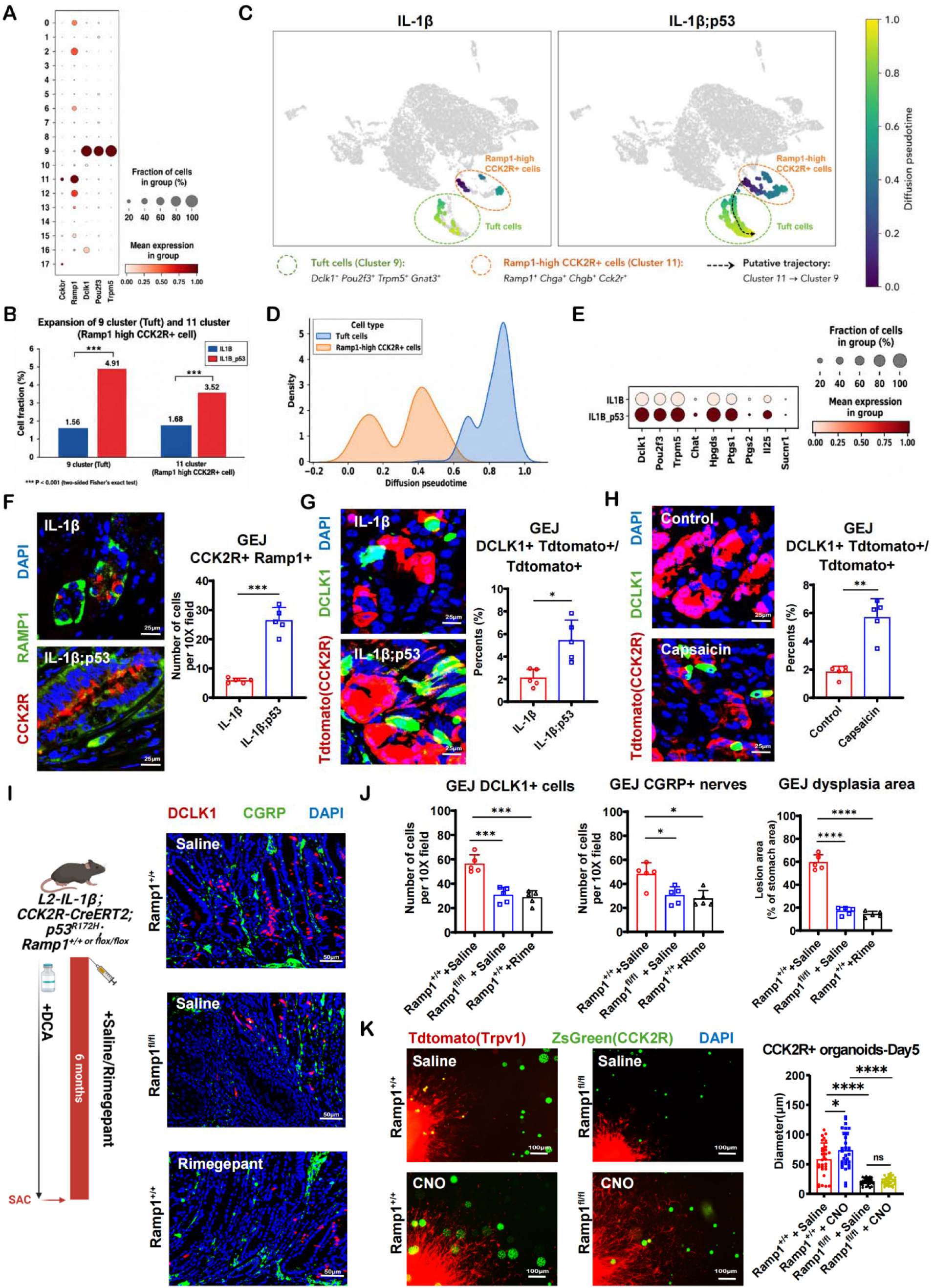
A neuro-responsive RAMP1-high CCK2R+ progenitor state mediates CGRP-dependent epithelial remodeling during GEJ tumorigenesis. (**A**) Dot plot of *Cck2r, Ramp1, Dclk1, Pou2f3,* and *Trpm5* expression across epithelial clusters from reanalysis of scRNA-seq data (GSE246280) from IL-1β and IL-1β; p53 GEJ epithelium; cluster 9, tuft cells; cluster 11, Ramp1-high CCK2R+ cells. (**B**) Fractions of cluster 9 (tuft) and cluster 11 (Ramp1-high CCK2R+) cells among epithelial cells in IL-1β versus IL-1β; p53 mice (two-sided Fisher’s exact test, \*\*\**P* < 0.001). (**C**) Diffusion-pseudotime maps of IL-1β and IL-1β; p53 epithelium highlighting tuft cells (cluster 9, green dashed outline) and Ramp1-high CCK2R+ cells (cluster 11, orange dashed outline); black dashed arrow indicates the putative trajectory from cluster 11 to cluster 9. (**D**) Density distributions of diffusion pseudotime for tuft cells and Ramp1-high CCK2R+ cells. (**E**) Dot plot of tuft-cell effector and receptor genes (*Dclk1, Pou2f3, Trpm5, Chat, Hpgds, Ptgs1, Ptgs2, Il25, Sucnr1*) by genotype. (**F**) IF of RAMP1 and CCK2R at the GEJ of IL-1β and IL-1β; p53 mice, with quantification of CCK2R+RAMP1+ cells per 10× field. n = 5 mice per group. (**G**) Lineage tracing in *L2-IL1B; Cck2r-CreERT2; R26-tdTomato* mice: IF of DCLK1 and tdTomato with quantification of DCLK1+tdTomato+ cells as a percentage of tdTomato+ cells. n = 5 mice per group. (**H**) Same lineage-tracing readout in mice fed a control or capsaicin-containing diet (0.01% capsaicin in AIN-93G, 6 months). n = 5 mice per group. (**I**) Experimental scheme: *L2-IL1B; Cck2r-CreERT2; p53^R172H^* mice with epithelial Ramp1 intact (*Ramp1^+/+^*) or deleted (*Ramp1^fl/fl^*), treated with DCA for 6 months, with saline or rimegepant (4 mg kg ¹ intraperitoneally, twice weekly); IF of DCLK1 and CGRP in the three groups. n = 5 mice per group. (**J**) Quantification of GEJ DCLK1+ cells, CGRP+ nerves, and dysplasia area for the experiment in (I). n = 5 mice per group. (**K**) Nodose ganglion (NG)– organoid coculture: NG from *Trpv1-Cre; R26-hM3Dq; R26-tdTomato* mice cocultured with CCK2R+ (ZsGreen+) organoids from *Cck2r-CreERT2; R26-ZsGreen; Ramp1^+/+^* or *Ramp1^fl/fl^* mice, treated with saline or CNO; representative images (tdTomato, red; ZsGreen, green) and day-5 organoid diameters. n = 30 organoids per condition. Scale bars: 25 μm (**F–H**), 50 μm (**I**), 100 μm (**K**). For all panels, values indicate mean ± SEM. \**P* < 0.05, \*\**P* < 0.01, \*\*\**P* < 0.001, \*\*\*\**P* < 0.0001; ns: not significant. Schematic in (**I**) created with BioRender.com.

Features of this neuroepithelial differentiation program were also observed in human disease. Analysis of scRNA-seq data from five patients with GEJ adenocarcinoma, encompassing tumor, adjacent, esophageal, and gastric tissues *(fig. S3A)*, identified a tuft-like epithelial population characterized by expression of POU2F3, TRPM5, and GNAT3 *(fig. S3B)*. Tuft-like cells accounted for a greater proportion of epithelial cells in tumors than in adjacent tissues in the pooled dataset (0.57% versus 0.26%), although this trend did not reach statistical significance in the patient-level paired analysis *(fig. S3, C and D)*. Similar to the findings in mouse tissues, pseudotime analysis placed the CCKBR+RAMP1+ epithelial population upstream of tuft-like cells, with progressive acquisition of the tuft-cell transcriptional program along the inferred differentiation trajectory *(fig. S3E)*, consistent with recent single-cell analyses linking GEJ epithelial states to metaplasia and cancer (38). Independent analyses of TCGA and GTEx datasets further showed increased expression of the tuft-cell markers POU2F3 and TRPM5 in esophageal and GEJ adenocarcinoma relative to normal tissues (32,39) *(fig. S3F)*. Together, these findings support a potentially important role for tuft-like cells in human GEJ tumorigenesis.

In vivo, CCK2R+RAMP1+ epithelial cells significantly increased in p53-mutant *L2-IL1B* mice relative to *L2-IL1B* controls. *(Fig. 2F)*. Lineage tracing in *L2-IL1B; Cck2r-CreERT2; R26-tdTomato* mice showed that p53-driven progression significantly increased the fraction of DCLK1+ tuft cells derived from CCK2R+ progenitors *(Fig. 2G)*, and chronic capsaicin-mediated activation of TRPV1-lineage sensory neurons (40) further enhanced tuft-cell differentiation from these progenitors *(Fig. 2H and fig. S3G)*, indicating that sensory neuronal activity promotes tuft-cell differentiation from neuro-responsive epithelial progenitors.

Epithelial CGRP–RAMP1 signaling was functionally required for this differentiation program. Conditional deletion of Ramp1 in CCK2R+ epithelial cells on the p53-mutant background, or pharmacological blockade of CGRP signaling with rimegepant—an orally available CGRP receptor antagonist approved for migraine (41)—markedly reduced DCLK1+ tuft cells, diminished CGRP+ sensory innervation, and suppressed dysplastic progression *(Fig. 2, I and J, and fig. S3H)*. Activation of TRPV1-lineage neurons robustly promoted the growth of control CCK2R+ organoids but failed to stimulate organoids lacking epithelial Ramp1 (*Fig. 2K and fig. S3I*), establishing epithelial RAMP1 as essential for sensory neuron–induced progenitor expansion and tuft-cell differentiation. These results extend the CGRP–RAMP1 axis recently implicated in established gastric cancer (30) to the earliest stages of the metaplasia-to-dysplasia transition at the GEJ.

### A bile acid-supported *P. melaninogenica* niche drives sensory neuroepithelial remodeling during GEJ tumorigenesis

We next sought the upstream environmental cues responsible for sensory nerve remodeling. Because the microbiota was a candidate driver, we compared conventionally housed and germ-free *L2-IL1B* mice (42). Germ-free conditions markedly reduced CGRP+ sensory innervation without affecting DCLK1+ tuft-cell abundance *(Fig. 3A)*, indicating that the microbiota selectively regulates sensory nerve remodeling rather than tuft-cell numbers per se.

**Fig. 3.**
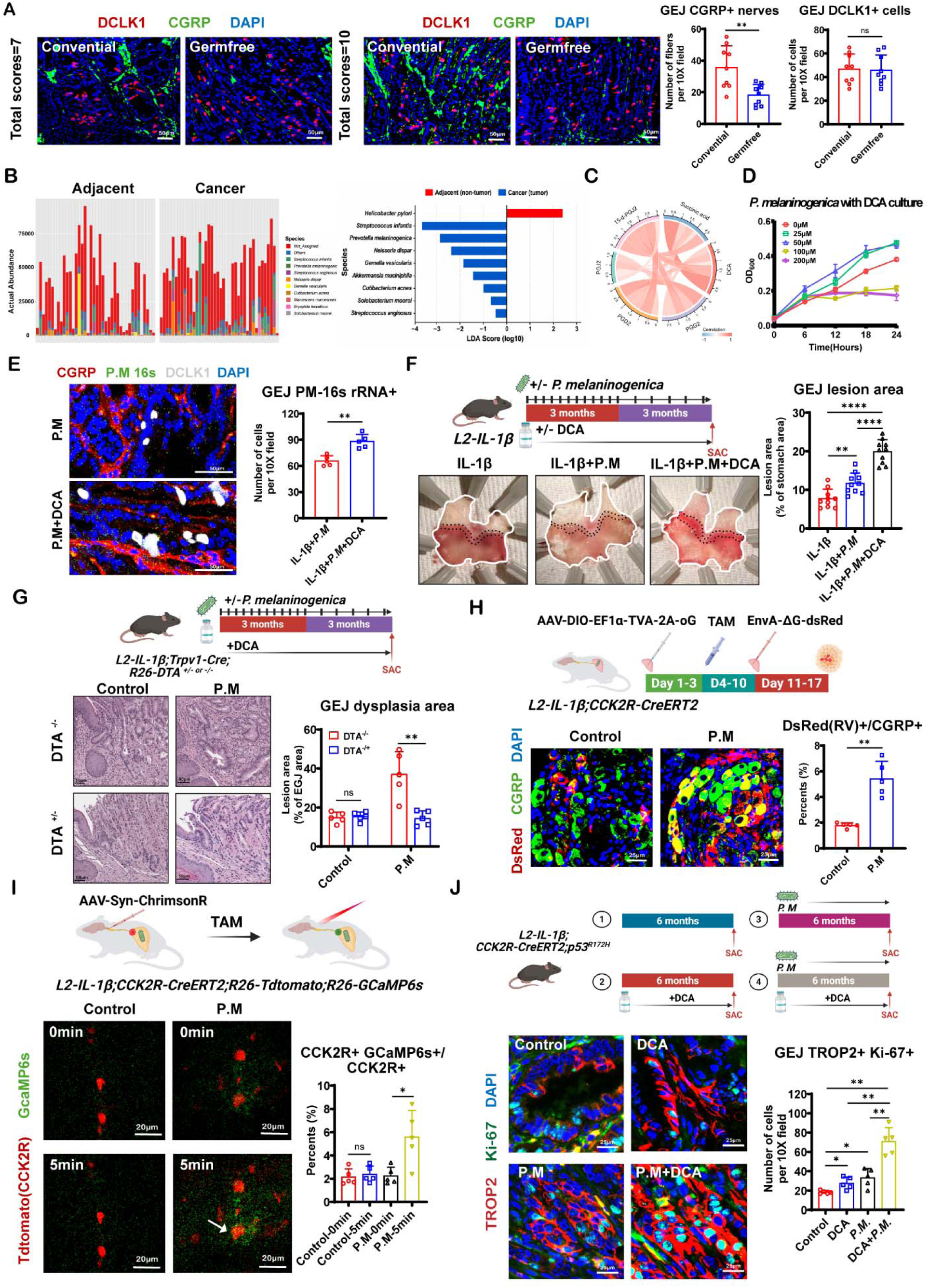
A bile acid-supported *P. melaninogenica* niche drives sensory neuroepithelial remodeling during GEJ tumorigenesis. (**A**) IF of DCLK1 and CGRP at the GEJ of conventionally housed and germ-free *L2-IL1B* mice at matched total dysplasia scores (score 7 and score 10), with quantification of CGRP+ nerves and DCLK1+ cells. n = 9 mice per group. (**B**) 16S rRNA sequencing of paired adjacent and tumor tissues from 40 patients with GEJ adenocarcinoma: species-level relative-abundance stacked bars (left) and LEfSe linear discriminant analysis (LDA) scores (right). (**C**) Circos plot of correlations between tumor-enriched bacterial species and tumor-enriched metabolites (including succinic acid, deoxycholic acid, and 15-d-PGJ2) in the same cohort. (**D**) Growth of *P. melaninogenica* (OD600 over 24 h) in medium containing 0–200 μM DCA. (**E**) RNAscope in situ hybridization for *P. melaninogenica* 16S rRNA combined with CGRP and DCLK1 IF at the GEJ of colonized mice with or without DCA, with quantification of *P. melaninogenica*-16S rRNA+ cells per 10× field. n = 5 mice per group. (**F**) Colonization scheme (*L2-IL1B* mice gavaged with *P. melaninogenic*a, with or without DCA, over 6 months), representative whole-mount stomachs, and quantification of GEJ lesion area. n = 10 mice per group. (**G**) Sensory-neuron ablation: *L2-IL1B; Trpv1-Cre; R26-DTA* mice (*DTA^+/−^*, ablated; *DTA^−/−^*, control) colonized with or without *P. melaninogenica* plus DCA; H&E sections and quantification of GEJ dysplasia area. n = 5 mice per group. (**H**) Monosynaptic retrograde rabies tracing from CCK2R+ epithelial cells (AAV-DIO-EF1a-TVA-2A-oG helper followed by EnvA-ΔG-dsRed rabies virus after tamoxifen induction) in control versus *P. melaninogenica*–colonized *L2-IL1B; Cck2r-CreERT2* mice; IF of dsRed (rabies virus) and CGRP in NG with quantification of dsRed(RV)+/CGRP+ neurons. n = 5 mice per group. (**I**) In vivo calcium imaging of CCK2R+ epithelial cells (*L2-IL1B; Cck2r-CreERT2; R26-tdTomato; R26-GCaMP6s*) during optogenetic activation of vagal sensory afferents (AAV-Syn-ChrimsonR injected into the left nodose ganglion): GCaMP6s and tdTomato signals at 0 and 5 min after light stimulation in control versus colonized mice, with quantification of GCaMP6s+ cells among CCK2R+ cells. n = 5 mice per group. (**J**) Four-arm experiment (control, DCA, *P. melaninogenica*, DCA + *P. melaninogenica*; 6 months): IF of TROP2 and Ki-67 with quantification of GEJ TROP2+Ki-67+ cells. n = 5 mice per group. Scale bars: 50 μm (**A, E**), 20 μm (**B, H, I**), 25 μm (**J**). For all panels, values indicate mean ± SEM. \**P* < 0.05, \*\**P* < 0.01, \*\*\**P* < 0.001, \*\*\*\**P* < 0.0001; ns: not significant. Schematic in (**F-J**) created with BioRender.com.

To identify microbiota-derived factors associated with nerve remodeling in human disease, we performed paired 16S rRNA sequencing and untargeted metabolomic profiling on tumors and adjacent tissues from 40 patients with GEJ adenocarcinoma *(fig. S4A)*. Tumor tissues showed extensive microbial community remodeling, with increased relative abundance of Bacteroidetes and Firmicutes at the phylum level and of *Prevotella* and *Streptococcus* at the genus level *(fig. S4, B to G)*; enrichment of these taxa was validated in The Cancer Microbiome Atlas (TCMA) (43), in which *Prevotella* abundance correlated positively with neuronal development and neurite-related genes, including TUBB2B, FGF11, and FGF14 *(fig. S5, A to C)*. Linear discriminant analysis identified *Streptococcus infantis* and *Prevotella melaninogenica* (*P. melaninogenica*) among the species most enriched in tumors (*Fig. 3B*). Because *Streptococcus infantis* is generally considered an oral commensal, whereas *P. melaninogenica* is an oral pathobiont associated with inflammatory disease (44), we focused on *P. melaninogenica* to investigate its potential role in tumor progression. Untargeted metabolomics separated tumor from adjacent tissue in both ion modes *(fig. S6, A to C)* and revealed enrichment of bile acids, prostaglandins, and succinate-related metabolites with extensive metabolic connectivity *(Fig. 3C)*; pathway analysis implicated lipid, amino acid, and central carbon metabolism, and network analysis converged on the tricarboxylic acid cycle, bacterial invasion of epithelial cells, and neuronal signaling modules, including synaptic vesicle cycling and neuroactive ligand–receptor interactions *(fig. S6D and fig. S7, A and B)*. Integrated microbiome–metabolome network analysis linked *Prevotella* to DCA, succinate, and multiple prostaglandin metabolites *(fig. S8A)*, suggesting functional coupling between microbial expansion and host metabolic remodeling.

Succinate is a major metabolic product of *P. melaninogenica* and related *Prevotella* species (45–46), whereas DCA is an established reflux-associated component at the GEJ and has been closely linked to Barrett’s esophagus and esophageal adenocarcinoma in mouse models (31,33). We therefore asked whether the bile acid environment preferentially supports *P. melaninogenica*. In vitro, DCA at concentrations ≤50 μM promoted dose-dependent bacterial growth, whereas higher concentrations were inhibitory (*Fig. 3D*). This growth-promoting range is comparable in magnitude to total DCA species measured in human refluxate (47). In *L2-IL1B* mice, *P. melaninogenica* colonization markedly increased bacterial abundance within GEJ lesions, where bacteria localized in close spatial proximity to both DCLK1+ tuft cells and CGRP+ sensory nerve endings, and DCA administration further enhanced colonization *(Fig. 3E)*. These findings indicate that DCA exposure can promote a local microbial niche that favors *P. melaninogenica* at the GEJ.

Colonization of *L2-IL1B* mice with *P. melaninogenica* significantly accelerated GEJ lesion development, an effect further enhanced by DCA *(Fig. 3F)*, with expansion of TROP2+ dysplastic epithelial cells and CGRP+ nerve endings within lesions, along with increased numbers of TRPV1+CGRP+ sensory neurons in the vagal NG *(fig. S9, A and B)*. To determine whether sensory neurons are required for microbiota-driven tumor progression, we genetically ablated TRPV1-lineage sensory neurons in *L2-IL1B*; *Trpv1-Cre; R26-DTA* mice. Despite continuous *P. melaninogenica* colonization and DCA exposure, sensory neuron ablation markedly reduced dysplastic progression compared with littermate controls *(Fig. 3G)*, establishing TRPV1-lineage sensory neurons as essential mediators of microbiota-driven GEJ tumorigenesis.

We next asked whether microbial colonization enhances anatomical communication between sensory neurons and epithelial progenitors. Monosynaptic rabies-virus tracing initiated from CCK2R+ epithelial cells (48–49) labeled CGRP+ sensory neurons in NG, and *P. melaninogenica* colonization significantly increased the proportion of rabies-labeled CGRP+ neurons *(Fig. 3H)*, indicating strengthened neuroepithelial connectivity. To test whether these connections are functionally active, we combined acute optogenetic activation of ChrimsonR-expressing sensory neurons with in vivo calcium imaging of GCaMP6s-expressing CCK2R-lineage epithelial cells. Following 5 min of optical stimulation of the NG, CCK2R-lineage epithelial cells exhibited robust calcium responses, which were significantly greater in colonized mice than in controls (*Fig. 3I*). These findings demonstrate that microbial colonization enhances functional sensory neuron-to-epithelium communication within the GEJ niche.

Accordingly, combined *P. melaninogenica* and DCA treatment significantly increased DCLK1+ tuft-cell abundance beyond either treatment alone *(fig. S9C)*, and after acquisition of the p53 mutation, the combination further accelerated tumor progression (fig. S9D), accompanied by marked expansion of TROP2+Ki-67+ epithelial cells, as well as tuft cells and CGRP+ sensory innervation *(Fig. 3J and fig. S9E)*. Sensory neurons thus serve as the principal conduit through which microbial signals are transmitted to the epithelium, coupling microbial remodeling to epithelial progenitor activation and dysplastic progression. These observations prompted us to identify the specific microbial metabolite that converts microbial colonization into neuronal activation.

### *P. melaninogenica*-derived succinate initiates neuroepithelial feed-forward signaling during GEJ tumorigenesis

Conditioned medium from *P. melaninogenica* activated TRPV1-lineage vagal sensory neurons exclusively through its low-molecular-weight (<3 kDa) fraction *(fig. S10, A and B)*, implicating small metabolites rather than secreted proteins. Succinate accumulated abundantly in *P. melaninogenica*–conditioned medium *(fig. S10C)* and exogenous succinate directly activated vagal sensory neurons, increased the proportion of responsive neurons, and stimulated CGRP release from cultured NG *(fig. S10, D and E)*. In contrast, although *Streptococcus anginosus* was likewise enriched in GEJ tumor tissues, its conditioned medium showed no significant increase in succinate and failed to elicit significant activation of TRPV1-lineage vagal sensory neurons *(fig. S10F)*. Colonization with *P. melaninogenica* also increased the proportion of SUCNR1+CGRP+ neurons within the NG *(fig. S10G)*, indicating that microbial colonization establishes a succinate-responsive sensory neuronal state.

Exogenous succinate phenocopied microbial colonization in vivo, accelerating dysplastic progression, enhancing epithelial proliferation within dysplastic regions, and increasing both tuft-cell abundance and sensory innervation in *L2-IL1B; Cck2r-CreERT2; Ramp1^+/+^* mice. These effects were completely abolished in *L2-IL1B; Cck2r-CreERT2; Ramp1^fl/fl^* mice, in which Ramp1 was conditionally deleted in CCK2R-lineage epithelial cells (*Fig. 4, A and B, and fig. S10H*). However, AAV-mediated deletion of Sucnr1 in TRPV1-lineage vagal sensory neurons only partially attenuated *P. melaninogenica*–induced dysplastic progression *(fig. S10I)*, and conditioned medium failed to directly stimulate axonal outgrowth of TRPV1-lineage neurons *(fig. S10J)*. Direct activation of sensory neurons alone therefore cannot fully explain the enhanced innervation induced by microbial colonization, suggesting an additional succinate-responsive cellular target within the GEJ microenvironment.

**Fig. 4.**
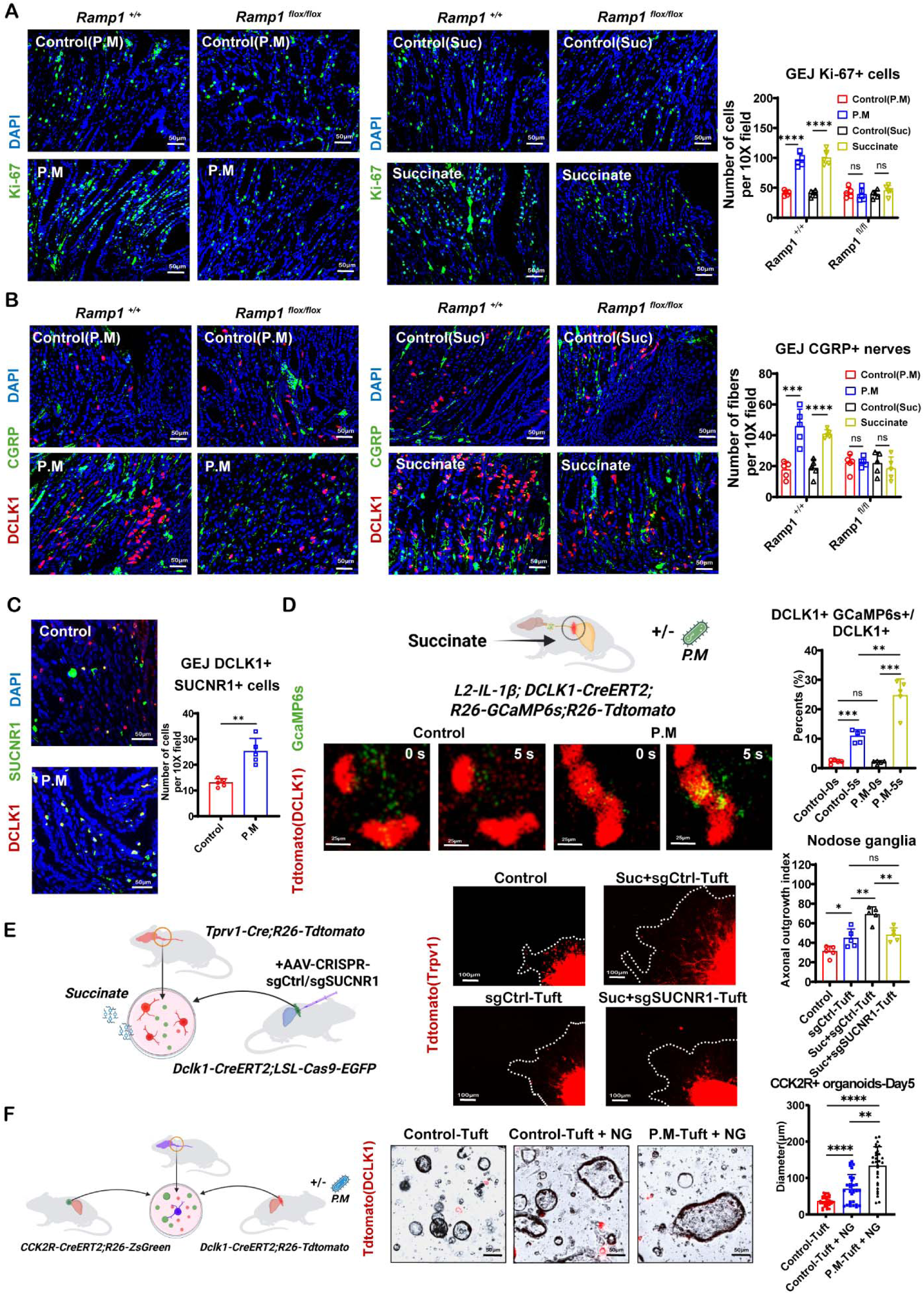
*P. melaninogenica*-derived succinate initiates neuroepithelial feed-forward signaling during GEJ tumorigenesis. (**A, B**) *L2-IL1B; Cck2r-CreERT2; Ramp1^+/+^* or *Ramp1^fl/fl^* mice treated with succinate (100 mM in drinking water) or *P. melaninogenica* colonization over 6 months, IF of Ki-67 and CGRP at the GEJ of *Ramp1^+/+^* mice and *Ramp1^fl/fl^*mice, with quantification of Ki-67+ cells and CGRP+ nerves. n = 5 mice per group. (**C**) IF of SUCNR1 and DCLK1 at the GEJ of control versus colonized mice, with quantification of SUCNR1+DCLK1+ cells. n = 5 mice per group. (**D**) In vivo calcium imaging of tuft cells (*L2-IL1B; Dclk1-CreERT2; R26-GCaMP6s; R26-tdTomato*) after luminal succinate administration, with or without *P. melaninogenica* colonization: GCaMP6s and tdTomato signals at 0 and 5 s, with quantification of GCaMP6s+ cells among DCLK1+ cells. n = 5 mice per group. (**E**) Neurite outgrowth from *Trpv1-Cre; R26-tdTomato* NG cocultured with tuft cells (*Dclk1-CreERT2; LSL-Cas9-EGFP*) transduced with AAV-CRISPR control (sgCtrl) or Sucnr1-targeting (sgSUCNR1) vectors, with or without succinate; representative images and axonal outgrowth index. n = 5 mice per group. (**F**) Coculture of CCK2R+ organoids (*Cck2r-CreERT2; R26-ZsGreen*) with tuft cells (*Dclk1-CreERT2; R26-tdTomato*) from control or colonized mice, with or without NG; representative bright-field/tdTomato images and day-5 organoid diameters. n = 30 organoids per condition. Scale bars: 50 μm (**B, C, F)**, 25 μm (**D**), 100 μm (**E**). For all panels, values indicate mean ± SEM. \**P* < 0.05, \*\**P* < 0.01, \*\*\**P* < 0.001, \*\*\*\**P* < 0.0001; ns: not significant. Schematic in (**D-F**) created with BioRender.com.

Intestinal tuft cells express SUCNR1 and respond to succinate (15–16, 50–51). In GEJ lesions, microbial colonization markedly increased the abundance of SUCNR1+ epithelial cells, which displayed a characteristic tuft-like morphology and predominantly co-expressed DCLK1 (*Fig. 4C*). In vivo calcium imaging in *L2-IL1B*; *Dclk1-CreERT2; R26-GCaMP6s* mice showed that luminal succinate administration elicited rapid calcium responses in DCLK1+ tuft cells, and these responses were potentiated in colonized mice *(Fig. 4D* and *Movie. S1)*. Succinate-activated tuft cells promoted sensory axonal outgrowth in co-culture—an effect abolished by CRISPR-mediated deletion of Sucnr1 in tuft cells *(Fig. 4E)*—and DCLK1+ tuft cells significantly enhanced the growth of CCK2R+ organoids in the presence of NG, an effect amplified in tuft cells isolated from colonized mice *(Fig. 4F)*. Succinate thus activates both vagal sensory neurons and SUCNR1-expressing tuft cells to initiate a neuroepithelial feed-forward circuit, implicating a tuft cell–derived mediator in its maintenance.

### Tuft cell-derived 15-d-PGJ2 sustains the neuroepithelial circuit during GEJ tumorigenesis

Although acetylcholine has been reported as a potential product of gastric tuft cells and may mediate rapid neuronal activation (25), ChAT, which encodes the acetylcholine-synthesizing enzyme choline acetyltransferase, showed low transcript abundance in tuft cells from mouse GEJ tumors *(Fig. 5A)*. We therefore reasoned that sustained sensory nerve remodeling during tumor progression might be driven by tuft cell–derived lipid mediators. Our human metabolomic analyses identified 15-deoxy-Δ12,14-prostaglandin J2 (15-d-PGJ2), the principal stable bioactive metabolite of the PGD2 pathway (52), among the most significantly enriched lipid metabolites in GEJ tumors, with abundance strongly associated with *P. melaninogenica* colonization *(fig. S8A)*. Tuft cells preferentially expressed Hpgds and Ptgs1, whereas Ptgs2 and Ptges expression remained minimal, indicating preferential activation of the PGD2 biosynthetic pathway rather than the canonical PGE2 pathway *(Fig. 5A and fig. S11A)*. Consistent with this profile, purified DCLK1+ tuft cells from *P. melaninogenica*–colonized mice secreted substantially more 15-d-PGJ2 than control tuft cells, whereas PGE2 secretion remained unchanged *(Fig. 5B and fig. S11B)*. DCLK1+HPGDS+ tuft cells expanded following colonization and further increased in p53-mutant *L2-IL1B* mice compared with *L2-IL1B* controls *(fig. S11C)*.

**Fig. 5.**
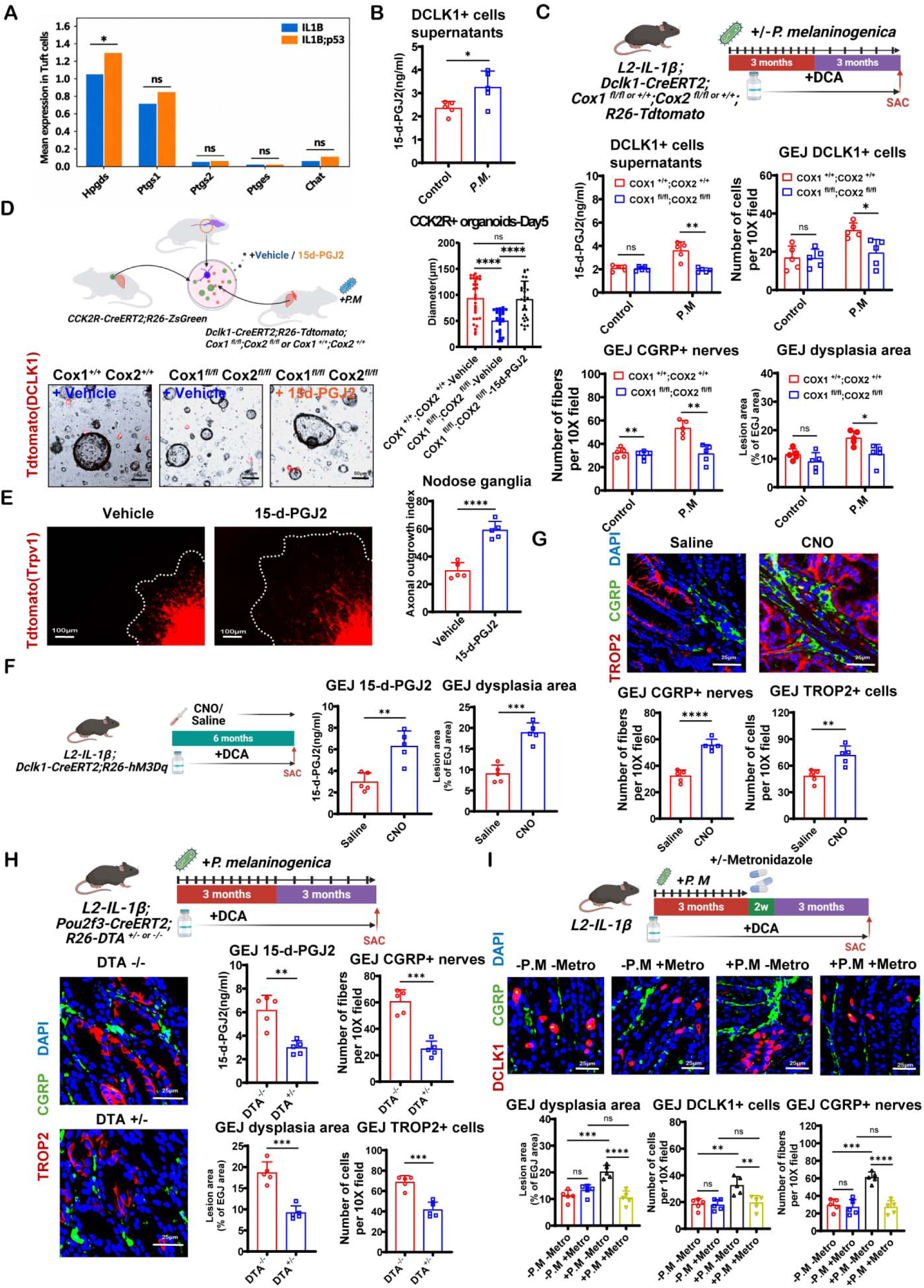
Tuft cell–derived 15-d-PGJ2 sustains the neuroepithelial circuit during GEJ tumorigenesis. (**A**) Mean expression of prostaglandin-pathway genes (*Hpgds, Ptgs1, Ptgs2, Ptges*) and *Chat* in tuft cells from IL-1β versus IL-1β; p53 GEJ epithelium (scRNA-seq). (**B**) 15-d-PGJ2 concentrations measured by ELISA in culture supernatants of DCLK1 tuft cells sorted from control or *P. melaninogenica*–colonized mice. n = 5 mice per group. (**C**) Tuft cell–specific prostaglandin-pathway deletion: *L2-IL1B; Dclk1-CreERT2; Cox1^fl/fl^* or *Cox1^+/+^*; *Cox2^fl/fl^*or *Cox2^+/+^*; *R26-tdTomato* mice colonized with *P. melaninogenica* plus DCA; quantification of 15-d-PGJ2 concentrations measured by ELISA in culture supernatants of DCLK1 tuft cells, GEJ DCLK1+ cells, CGRP+ nerves, and dysplasia area. n = 5 mice per group. (**D**) CCK2R+ organoid–tuft cell coculture with Cox1/Cox2-intact or –deficient tuft cells (from *P. melaninogenica*–colonized mice), with vehicle or 15-d-PGJ2 add-back; representative images and day-5 organoid diameters. n = 30 organoids per condition. (**E**) Neurite outgrowth from *Trpv1-Cre; R26-tdTomato* NG with vehicle or 15-d-PGJ2; axonal outgrowth index. n = 5 mice per group. (**F**) Chemogenetic activation of tuft cells: *L2-IL1B; Dclk1-CreERT2; R26-hM3Dq* mice treated with CNO or saline, with DCA, for 6 months; GEJ 15-d-PGJ2 concentrations and dysplasia area. n = 5 mice per group. (**G**) IF of CGRP and TROP2 in saline-versus CNO-treated mice, with quantification of CGRP+ nerves and TROP2+ cells. n = 5 mice per group. (**H**) Tuft-cell ablation: *L2-IL1B; Pou2f3-CreERT2; R26-DTA* mice (*DTA^+/−^* versus *DTA^−/−^*) colonized with *P. melaninogenica* plus DCA; IF of CGRP and TROP2, with quantification of 15-d-PGJ2, CGRP+ nerves, dysplasia area, and TROP2+ cells. n = 5 mice per group. (**I**) Bacterial depletion: *L2-IL1B* mice in a 2 × 2 design (± *P. melaninogenica* colonization; ± metronidazole, 7.1 mg kg ¹ twice daily by gavage for 14 days), with DCA; IF of DCLK1 and CGRP, with quantification of dysplasia area, DCLK1+ cells, and CGRP+ nerves. n = 5 mice per group. Scale bars: 50 μm (**D, G–I**), 100 μm (**E**). For all panels, values indicate mean ± SEM. \**P* < 0.05, \*\**P* < 0.01, \*\*\**P* < 0.001, \*\*\*\**P* < 0.0001; ns: not significant. Schematic in (**C, D, F, H, I**) created with BioRender.com.

To determine whether tuft cell–derived prostaglandins are required for neuroepithelial remodeling, we conditionally deleted Cox1/Cox2 specifically in DCLK1 tuft cells. Despite persistent *P. melaninogenica* colonization, disruption of prostaglandin biosynthesis markedly reduced tuft-cell production of 15-d-PGJ2, without significantly reducing PGE2 production *(Fig. 5C and fig. S11D)*. This deletion significantly attenuated dysplastic progression and reduced DCLK1 tuft-cell abundance and CGRP sensory innervation within lesions *(Fig. 5C)*. Loss of Cox1/Cox2 in tuft cells also impaired the ability of NG to support the growth of CCK2R+ organoids, whereas exogenous 15-d-PGJ2 completely rescued this defect *(Fig. 5D)*. Tuft cell– derived prostaglandin signaling is thus indispensable for maintaining neuroepithelial remodeling.

Unlike succinate, 15-d-PGJ2 alone failed to elicit overt neuronal calcium responses; however, pretreatment with 15-d-PGJ2 markedly enhanced neuronal responsiveness to subsequent succinate stimulation, significantly increasing the proportion of activated vagal sensory neurons *(fig. S11E)*. In addition, 15-d-PGJ2 directly promoted sensory axonal outgrowth in cultured NG (*Fig. 5E)*. Tuft cell–derived 15-d-PGJ2 therefore functions as a neuronal sensitizer that promotes sustained sensory nerve remodeling rather than acting as a fast neurotransmitter.

Chemogenetic activation of DCLK1+ tuft cells in *L2-IL1B*; *Dclk1-CreERT2; R26-hM3Dq* mice significantly increased intralesional 15-d-PGJ2 levels, concomitantly accelerated lesion growth and dysplastic progression, and expanded CGRP+ sensory nerve endings and TROP2+ dysplastic epithelial cells *(Fig. 5, F and G, and fig. S11F)*. Conversely, genetic ablation of tuft cells in *L2-IL1B*; *Pou2f3-CreERT2; R26-DTA* mice—targeting the POU2F3-dependent tuft-cell lineage (17, 53)—markedly reduced intralesional 15-d-PGJ2, sensory innervation, TROP2+ epithelial expansion, and dysplastic progression despite persistent colonization and DCA exposure *(Fig. 5H and fig. S11G)*. Finally, metronidazole-mediated depletion of *P. melaninogenica* simultaneously reduced GEJ tuft-cell abundance, sensory innervation, and dysplastic progression within GEJ lesions *(Fig. 5I and fig. S11, H and I)*. Microbial metabolites thus initiate activation of sensory neurons and tuft cells, whereas tuft cell–derived 15-d-PGJ2 perpetuates sensory nerve remodeling to maintain CGRP–RAMP1 signaling, driving sustained expansion of CCK2R+ progenitors and ultimately promoting GEJ tumorigenesis.

## DISCUSSION

This study defines a microbiota-driven neuroepithelial feed-forward circuit that converts environmental dysbiosis into persistent oncogenic signaling at the GEJ. Rather than acting as independent compartments, microbial metabolites, tuft cells, sensory neurons, and epithelial progenitors cooperate to establish a self-reinforcing signaling network that progressively remodels the premalignant niche. This framework extends current models of Barrett-associated tumorigenesis by identifying neuroepithelial communication as a central mechanism through which environmental information is translated into sustained epithelial remodeling and dysplastic progression.

Bile reflux is an established driver of Barrett’s esophagus and GEJ adenocarcinoma, and its carcinogenicity has been attributed largely to direct genotoxic injury of epithelial cells (12–13) and, more recently, to bile acid receptor signaling (54). Our findings provide, to our knowledge, the first evidence that bile acids also promote carcinogenesis indirectly—by reshaping microbial colonization. Low, reflux-compatible DCA concentrations selectively supported *P. melaninogenica*, whose metabolic output, rather than its biomass or a genotoxin, initiated oncogenic signaling. Microbial metabolites are increasingly recognized as cancer drivers, exemplified by colibactin, which alkylates epithelial DNA in colorectal cancer (4). The mechanism described here is distinct in kind: the microbial product acts not on epithelial DNA but on the tissue’s sensory apparatus—neurons and chemosensory tuft cells—converting a transient microbial signal into durable neural and epithelial remodeling.

This mechanism also contrasts with other bacteria implicated in upper gastrointestinal cancer. *Streptococcus anginosus* promotes gastric tumorigenesis through direct bacterial adhesion to epithelial cells via the TMPC–ANXA2 interaction and downstream MAPK signaling (55), and our human microbiome analyses likewise detected *Streptococcus* enrichment within GEJ tumors. However, conditioned medium from *Streptococcus anginosus* did not contain elevated succinate and failed to robustly activate TRPV1-lineage vagal sensory neurons, supporting a distinct, non-neurotropic mode of tumor promotion. *P. melaninogenica*, by contrast, emerged as the species most closely associated with bile acid remodeling, microbial metabolites, and neuronal signaling pathways, and it acts at a distance—through succinate—on neurons and tuft cells rather than through direct epithelial contact. Distinct members of the dysbiotic community may thus engage epithelial transformation through fundamentally different routes, with *P. melaninogenica* occupying the central microbial hub linking environmental dysbiosis to neuroepithelial remodeling.

Neural circuits are now recognized drivers of solid tumors, but neurons have generally been viewed as downstream responders to tissue injury or tumor-derived signals, and the cues that activate tumor-promoting nerves have remained unclear (22–23). Our data position vagal sensory neurons among the earliest environmental sensors in tumorigenesis and identify a microbial metabolite as one such activating signal. Sensory neurons are known to directly detect bacterial products, regulating pain, host defense, and meningeal immunity (18–19, 56), and to sense metabolites that govern host nutrition, physiology, and feeding (20–21); our findings extend this sensory modality to cancer initiation. The principal epithelial target of this neural activity is a transient RAMP1-high CCK2R+ progenitor state that subsequently differentiates toward the tuft-cell lineage, linking neural activity to epithelial lineage plasticity rather than to the growth of mature epithelium. This progenitor-centered view complements the CGRP–RAMP1 axis described between nociceptors and established gastric cancer cells (30) and may explain how neural activity becomes coupled to expansion of a metaplastic and dysplastic lineage. Additionally, in the small intestine, activation of nociceptive nerves can promote progenitor cell proliferation and tuft cell differentiation/activation through the CGRP receptor to regulate type 2 immunity (57).

Direct chemical communication between epithelial chemosensory cells and sensory neurons is well established for enteroendocrine cells, which form putative synapses with vagal neurons to transduce nutrient signals (20–21), and tuft cell–neuron communication has been proposed at mucosal barriers (17). Our findings extend this biology in two ways. First, tuft cells are not merely environmental sensors: through 15-d-PGJ2–mediated neuronal sensitization, they actively sustain nerve remodeling during tumorigenesis. Second, the circuit divides labor between metabolites—microbiota-derived succinate initiates activation of both neurons and tuft cells, whereas tuft cell–derived 15-d-PGJ2 maintains and amplifies the signal—providing a mechanistic explanation for how transient environmental perturbations are converted into persistent neuroepithelial remodeling during tumor initiation.

These findings have potential therapeutic implications. Multiple components of the circuit— microbial communities, bile acid metabolism, succinate signaling, tuft-cell prostaglandin biosynthesis, and CGRP–RAMP1 signaling—are potentially amenable to therapeutic intervention. In particular, the ability of the clinically approved CGRP receptor antagonist rimegepant (22, 41) to phenocopy genetic disruption of epithelial Ramp1 highlights neural signaling as a clinically actionable vulnerability during early tumorigenesis. Our results also offer a mechanistic explanation for the established epidemiological association between periodontal disease and increased risks of esophageal and gastric adenocarcinoma (10–11): the oral commensal *P. melaninogenica* emerges as a central driver of neuroepithelial remodeling in GEJ tumors, suggesting that maintaining oral health and selectively reducing pathogenic oral bacteria may represent an upstream strategy for cancer interception, alongside neutralization of toxic bile acids or inhibition of PGD2–15-d-PGJ2 signaling. Notably, targeting downstream pathways of the microbial driver—such as succinate–SUCNR1 signaling, tuft-cell prostaglandin biosynthesis, or CGRP–RAMP1 signaling—may prove more effective and feasible than attempting to eradicate the microbes themselves, which are oral commensals embedded within complex communities and prone to recolonization.

Several limitations should be acknowledged. Our human microbiome and metabolomic analyses are associative, and direct causal validation in patients remains challenging. In addition, although *P. melaninogenica* emerged as the principal microbial hub, other microbial species are likely to cooperate within the GEJ ecosystem and warrant further investigation. Barrett’s esophagus and esophageal adenocarcinoma are widely believed to arise from the combined actions of acid and bile (1–3), whereas our models do not recapitulate acid reflux; notably, both sensory neurons and tuft cells can sense pH, and acid exposure could in principle engage neural signaling as well (58–59). Moreover, although tuft cells expand early during dysplasia, their numbers decline in high-grade dysplasia and established adenocarcinoma (60), suggesting that tuft cell–neuron signaling is most important during early tumor initiation rather than late-stage progression. Finally, our study focused on peripheral vagal sensory neurons; whether central neural circuits also regulate or amplify this feed-forward network requires further investigation.

## Acknowledgments

We acknowledge support from Columbia University shared resources and thank Dajiang “Kevin” Sun and Tingting Sun (Molecular Pathology/MPSR, Columbia University) for their expertise and assistance during this project. We thank Dr. Theresa Swayne and Dr. Yanyan Chen from Columbia University for their contributions to fluorescence imaging of live animal cells. We also thank Dr. Yi Wang from Southwest University and Dr. Gang Ma from Tianjin Medical University for their contributions to the analysis of human samples. We thank Bertram Bengsch for support within the The Freiburg Mass Cytometry Unit.

## Author contributions

Conceptualization: T.C.W., Y.Z., J.A.A., M.Q., J.G.F.

Methodology: Y.Z., Z.Y., F.W., P.Z., X.Z., M.B., J.Q., S.L., Q.T.W., R.T., M.H., J.L., H.K., J. A.A, Y.O., H.Z., B.Z., N.H., M.Q.

Investigation: Y.Z., Z.Y., M.B., M.Q.

Visualization: Y.Z., Z.Y., F.W., P.Z.

Funding acquisition: T.C.W., M.Q.

Project administration: T.C.W., L.B.Z., K.L.V.

Supervision: T.C.W., J.G.F., J. A.A.

Writing – original draft: Y.Z.

Writing – review & editing: T.C.W., Y.Z., J.A.A., Z.Y., H.K

## Data and materials availability

The human single-cell RNA sequencing dataset generated in this study has been deposited in the National Genomics Data Center (NGDC, China National Center for Bioinformation) under accession number HRA017684. Mouse GEJ epithelial single-cell RNA sequencing data were obtained from the Gene Expression Omnibus (GEO) under accession number GSE246280. Human microbiome data were obtained from The Cancer Microbiome Atlas (TCMA; https://tcma.pratt.duke.edu/). Public transcriptomic data from The Cancer Genome Atlas (TCGA) and the Genotype-Tissue Expression (GTEx) project were used for validation analyses. The imaging mass cytometry (IMC) dataset was generated by the laboratory of Michael Quante in the The Freiburg Mass Cytometry Unit . Reagents are available on request from the lead contact with a completed materials transfer agreement.

## Supplementary Materials

### Materials and Methods

Figs. S1 to S11

Movie S1

