## Supplemental Methods and Figures for "A microbiota-driven neuroepithelial feed-forward circuit drives gastroesophageal junction tumorigenesis"

**Materials and Methods**

**Animals**

All animal experiments were conducted in accordance with National Institutes of Health guidelines and were approved by the Institutional Animal Care and Use Committee (IACUC) of Columbia University, Massachusetts Institute of Technology and Klinikum Rechts der Isar, TUM. Mice were maintained under specific pathogen-free conditions on a 12-h light–dark cycle (lights on, 07:00–19:00) at 22 °C and 40–50% humidity, with ad libitum access to food and water. Unless otherwise specified, experiments were performed in 8–12-week-old mice. Both male and female mice were used and age- and sex-matched across experimental groups. Littermate controls were used when possible.

*Dclk1-CreERT2, Cck2r-CreERT2,* and *L2-IL1B* mice have been described previously. The following mouse strains were obtained from The Jackson Laboratory: C57BL/6J (stock no. 000664; wild-type, WT), *B6.129P2-Gt (ROSA)26Sortm1(DTA)Lky/J* (stock no. 009669; *R26-DTA*), *B6. Cg-Gt (ROSA)26Sortm9(CAG-tdTomato) Hze/J* (stock no. 007909; *R26-tdTomato*), *B6.129-Trpv1tm1(cre)Bbm/J* (stock no. 017769; *Trpv1-Cre*), *B6N;129-Tg (CAG-CHRM3*, -mCitrine)1Ute/J* (stock no. 026220; *R26-hM3Dq*), and *B6J.Cg-Gt (ROSA)26Sortm96(CAG-GCaMPs) Hzee/MwarJ* (stock no. 028866; *R26-GCaMP6s*). *Ramp1^fl/fl^* mice were provided by Dr. Isaac Chiu (Harvard Medical School). *Trp53LSL-R172H* mice were provided by Dr. Kenneth Olive (Columbia University). *Pou2f3-CreERT2* mice were provided by Dr. Jakob von Moltke (University of Washington). *Ptgs1^fl/fl^* and *Ptgs2^fl/fl^* mice were provided by Dr. Harvey Herschman (University of California, Los Angeles). All strains were maintained on a C57BL/6 background and crossed to generate the experimental genotypes described in the text, including *L2-IL1B; Cck2r-CreERT2; LSL-p53^R172H+/-^*, *L2-IL1B; Trpv1-Cre; R26-hM3Dq*, *L2-IL1B; Trpv1-Cre; R26-DTA*, *L2-IL1B; Cck2r-CreERT2; R26-tdTomato*, *L2-IL1B; Cck2r-CreERT2; Ramp1^fl/fl^; LSL-p53^R172H+/-^*, *Dclk1-CreERT2; R26-hM3Dq*, *Dclk1-CreERT2; R26-GCaMP6s*, *Dclk1-CreERT2; Ptgs1^fl/fl^; Ptgs2^fl/fl^*, and *Pou2f3-CreERT2; R26-DTA* mice.

For induction of CreERT2-mediated recombination, tamoxifen (Sigma) was administered at 100 mg kg⁻¹ unless otherwise specified. Sample sizes were based on prior experience with these models and were sufficient to detect biologically meaningful differences. No statistical methods were used to predetermine sample size. Animals were randomly assigned to experimental groups when possible. Investigators were not blinded during experiments.

Germ-free (GF) mice were generated by embryo transfer rederivation of conventional L2-IL1B mice into GF Swiss recipient mothers17 and maintained in sterile isolators on autoclaved food (RMH3000; Purina, St. Louis, MO) comparable to the standard chow and water. Sterility was confirmed every other week by 16S polymerase chain reaction; aerobic, anaerobic and fungal culture; and Gram stains of fecal smears. Age-matched conventionally housed L2-IL1B mice served as controls.

**Mouse models of GEJ tumorigenesis and in vivo treatments**

For dietary intervention studies conducted in Germany, mice were randomly assigned to treatment groups at 8–9 weeks of age and fed either a high-fat diet (HFD; Ssniff, S5745-E712) or a matched control diet (CD; Ssniff, S5745-E702).

For chemogenetic activation of TRPV1-lineage sensory neurons, *L2-IL1B; Trpv1-Cre; R26-hM3Dq* mice received clozapine-N-oxide (CNO; Tocris) by oral gavage at 0.3 mg kg⁻¹ three times per week for the durations indicated. Control mice received vehicle by oral gavage on the same schedule.

For chemogenetic activation of Dclk1-lineage tuft cells, *Dclk1-CreERT2; L2-IL1B; R26-hM3Dq* mice were first treated with tamoxifen to induce Cre-dependent hM3Dq expression. Following tamoxifen induction, mice received CNO by oral gavage at 0.3 mg kg⁻¹ three times per week for the durations indicated. Control mice received vehicle by oral gavage on the same schedule.

For chronic dietary capsaicin exposure, mice were fed an AIN-93G-based diet supplemented with 0.01% capsaicin (Dyets, Inc.) for 6 months. Control mice received the AIN-93G diet without capsaicin.

To inhibit CGRP signaling during tumor progression, rimegepant (MedChemExpress, HY-15498) was administered intraperitoneally at 4 mg kg⁻¹ twice weekly for 6 months. Control mice received vehicle injections on the same schedule.

Deoxycholic acid (DCA; Sigma-Aldrich) was administered in drinking water at 0.2% (w/v) for the durations indicated as previously described (31,33). Sodium succinate (Sigma-Aldrich) was administered in drinking water at 100 mM for 6 months. Control mice received vehicle-matched drinking water.

**Bacterial strains, culture, and conditioned medium**

*Prevotella melaninogenica* (ATCC 25845) and *Streptococcus anginosus* (ATCC 33397) were cultured anaerobically (85% N₂, 10% H₂, 5% CO₂) at 37 °C in brain heart infusion broth supplemented with hemin (5 μg ml⁻¹) and vitamin K1 (1 μg ml⁻¹) / Gifu anaerobic medium. Growth was monitored by optical density at 600 nm (OD600).

For DCA dose–response experiments, *P. melaninogenica* was inoculated into medium containing deoxycholic acid at final concentrations of 0, 10, 25, 50, 100, and 200 μM, and growth was recorded by OD600 measurements every 2 h for 24–48 h. All assays were performed in at least three independent biological replicates.

Conditioned medium (CM) was prepared from late-log-phase cultures (OD600 ≈ 0.8), clarified by centrifugation (5,000 × g, 15 min, 4 °C), and sterilized through 0.22-μm filters. Low-molecular-weight fractions (<3 kDa) were obtained by ultrafiltration using Amicon Ultra centrifugal filters (3 kDa molecular-weight cutoff, Millipore) according to the manufacturer's instructions; the flow-through was collected and used for stimulation experiments. Succinate concentrations in CM and fractions were quantified using a succinate assay kit (Enzo life science).

**Bacterial colonization and depletion in mice**

For colonization experiments, 8–12-week-old L2-IL1B mice received *P. melaninogenica* by oral gavage (1 × 10⁹ CFU in 200 μl anaerobic PBS) once weekly for the first 3 months and once every 2 weeks for the subsequent 3 months. Control mice received vehicle by oral gavage on the same schedule. Where indicated, DCA was administered concomitantly as described above. Colonization in GEJ tissues was assessed by RNAscope in situ hybridization using a probe specific for P. melaninogenica 16S rRNA, as described below.

For bacterial depletion, metronidazole (Sigma-Aldrich) was administered by oral gavage at 7.1 mg kg⁻¹ per dose twice daily (total daily dose, 14.2 mg kg⁻¹) for 14 consecutive days. The efficacy of depletion was assessed by RNAscope in situ hybridization using the same P. melaninogenica-specific 16S rRNA probe.

**Human specimens**

Paired tumor and adjacent non-tumor tissues were collected from 40 patients with gastroesophageal junction (GEJ) adenocarcinoma undergoing surgical resection at Fujian Cancer Hospital between 2023 and 2024. For five of these patients, matched samples from four anatomical sites were collected: tumor tissue, tumor-adjacent gastric tissue, tumor-adjacent esophageal tissue, and distant gastric tissue located more than 10 cm from the tumor. The study was approved by the Institutional Review Board of Fujian Cancer Hospital (approval No. K2023-463-01), and written informed consent was obtained from all participants. Patients who had received neoadjuvant therapy, antibiotic treatment within 4 weeks before surgery, or proton pump inhibitor treatment within 2 weeks before surgery were excluded. Tissues were collected within 30 min of resection. Separate tissue portions were snap-frozen in liquid nitrogen for nucleic acid and metabolite extraction or fixed in formalin and embedded in paraffin for histological analysis.

**16S rRNA gene sequencing and analysis**

Microbial DNA extraction, 16S rRNA gene sequencing, and bioinformatic analysis were performed by Novogene (China). DNA was extracted from frozen tissue samples and assessed for concentration and quality before amplification. The V3–V4 region of the bacterial 16S rRNA gene was amplified using primers 341F (5′-CCTAYGGGRBGCASCAG-3′) and 806R (5′-GGACTACNNGGGTATCTAAT-3′). Amplicons were purified, pooled, and used for library preparation, followed by paired-end sequencing on an Illumina platform. Raw reads underwent quality filtering and chimera removal using the Novogene analysis pipeline. Taxonomic annotation was performed against a reference 16S rRNA gene database. Microbial community composition and alpha and beta diversity were analyzed using the resulting feature abundance table. Differences in community structure were evaluated by permutational multivariate analysis of variance (PERMANOVA), and differentially abundant taxa were identified using linear discriminant analysis effect size (LEfSe).

**Untargeted metabolomics**

Untargeted metabolomic profiling of frozen tissue samples was performed by Novogene (China) using liquid chromatography–tandem mass spectrometry (LC–MS/MS). Tissue samples were homogenized and subjected to metabolite extraction according to the service provider’s protocol. Extracts were clarified before analysis by ultra-high-performance liquid chromatography coupled to high-resolution mass spectrometry in positive and negative electrospray ionization modes. Raw data were processed for peak detection, retention-time alignment, and peak integration. Metabolites were annotated by matching accurate mass and MS/MS fragmentation spectra against reference databases. Processed peak intensities were used to assess relative metabolite abundances.

**Public dataset analyses**

Microbial abundance data for all available bacterial taxa in the esophageal and gastric adenocarcinoma cohorts were obtained from The Cancer Microbiome Atlas (TCMA; https://tcma.pratt.duke.edu). Associations between bacterial abundance and host gene expression were evaluated across these cohorts.

Gene expression data from The Cancer Genome Atlas (TCGA) esophageal adenocarcinoma and stomach adenocarcinoma cohorts and normal esophageal and gastric tissues from the Genotype-Tissue Expression (GTEx) project were analyzed using the Xiantao online analysis platform(www.xiantaozi.com).

**Single-cell RNA sequencing analysis**

Mouse GEJ epithelial single-cell RNA sequencing (scRNA-seq) data from *L2-IL1B* and *L2-IL1B; Cck2r-CreERT2; p53^R172H^* mice were reanalyzed from our previously published dataset (GEO accession No. GSE246280). Human scRNA-seq data from paired GEJ adenocarcinoma and adjacent non-tumor tissues were generated in this study and deposited in the Genome Sequence Archive for Human (GSA-Human) at the National Genomics Data Center (NGDC; accession No. HRA017684).

Computational analyses were performed in Python. Raw gene–cell count matrices were subjected to quality control, excluding cells with fewer than 200 or more than 6,000 detected genes, or with more than 10% of counts assigned to mitochondrial genes. Gene expression counts were normalized and log-transformed, and highly variable genes were selected for dimensionality reduction by principal component analysis (PCA). Uniform manifold approximation and projection (UMAP) was used to visualize cell populations, and clusters were annotated using canonical marker genes. Differential gene expression and trajectory analyses were performed to characterize transcriptional differences and relationships among epithelial cell populations. For pseudotime analysis, the RAMP1-high CCK2R⁺ progenitor population was designated as the starting population.

**Imaging mass cytometry**

Formalin-fixed, paraffin-embedded (FFPE) GEJ sections (4 μm) were stained with a panel of metal-conjugated antibodies targeting DCLK1, TUBB3 (βIII-tubulin), CGRP, EpCAM, TROP2, CD45, CD31, αSMA, Ki-67, and additional lineage markers using Maxpar reagents (Standard BioTools). Regions of interest (ROIs) encompassing the squamocolumnar junction (SCJ) and metaplastic and dysplastic lesions were selected using corresponding H&E-stained serial sections as a guide. Data were acquired on a Hyperion Imaging System (Standard BioTools) at a laser ablation frequency of 200 Hz and a spatial sampling resolution of 1 μm per pixel. Single-cell segmentation was performed using ilastik and CellProfiler within the steinbock workflow, incorporating nuclear signals from the 191Ir/193Ir DNA intercalator and membrane-marker channels. Cell phenotypes were assigned by PhenoGraph clustering followed by expert review of marker expression.

**Spatial analysis of tuft cells and β3-tubulin–positive neural structures**

IMC data were processed using the established image-analysis workflow for highly multiplexed tissue imaging, including image generation, cell segmentation, and phenotypic annotation. Segmentation was based on nuclear and cellular boundaries and therefore captured neuronal cell bodies more reliably than thin axonal or neurite processes; cells annotated as tuft cells or neurons were retained as the segmented tuft-cell and neuron-like populations, respectively. To assess neural structures independently of cell segmentation, the β3-tubulin channel was additionally analyzed as a continuous image field.

β3-tubulin images were Gaussian-smoothed (σ = 1 pixel) without additional background subtraction. For within-ROI localization and distance analyses, β3-tubulin–bright pixels were defined using the ROI-specific 95th percentile of positive tissue signal. For comparisons of β3-tubulin signal burden between ROIs, a variable-area high-signal mask was defined as the median positive β3-tubulin intensity plus four median absolute deviations. Connected high-signal objects smaller than five pixels were excluded in both approaches to reduce isolated pixel noise.

Mouse-level tuft-cell densities and β3-tubulin high-signal area fractions were summarized across sampled ROIs and related by Spearman rank correlation. Area-normalized tuft-cell counts within 10, 25, and 50 μm of β3-tubulin–bright pixels were compared across dysplasia scores, and β3-tubulin–bright pixel area was quantified within the whole ROI and within annotated lesion, epithelium, and stroma compartments. To test whether tuft cells were specifically enriched near neural structures, tuft cells were grouped by ROI and local epithelial tissue compartment, and their median nearest distance to stromal β3-tubulin–bright pixels was compared with a null distribution generated from 1,000 random permutations of an equal number of non-tuft epithelial cells from the same ROI. ROI contrasts were aggregated within each mouse weighted by tuft-cell number, and the mouse-level directional contrast was tested against zero using a one-sided Wilcoxon signed-rank test.

Directional tuft-cell→neuron spatial preference was independently quantified using COZIpy (v0.1.4) with a 20-μm neighborhood radius and 1,000 within-ROI label permutations. COZI z-scores were normalized by the square root of the total number of cells in each ROI to reduce dependence on ROI cellularity. Formal progression models required both cell types to be represented by at least 20 cells per qualifying ROI and pair support in at least 12 ROIs and 8 mice spanning all three aggregated dysplasia scores. Associations with dysplasia were tested using a linear mixed-effects model of the form standardized COZI score ~ standardized dysplasia + (1 | mouse). P values were adjusted across all tested directional cell-type pairs using the Benjamini–Hochberg method. Mouse identity was treated as the experimental unit throughout; repeated ROIs were either summarized at the mouse level or retained in mixed-effects models with a mouse-level random intercept.

**Immunofluorescence and histology**

GEJ tissues were fixed in 4% paraformaldehyde or 10% neutral-buffered formalin at 4 °C and embedded in paraffin or OCT compound. Tissue sections were prepared using standard histological procedures. For immunostaining, sections underwent antigen retrieval, were blocked with 10% BSA for 1 h at room temperature, and incubated with primary antibodies overnight at 4 °C. Primary antibodies included anti-TRPV1 (1:200, Alomone Labs, ACC-030), anti-CGRP (1:200, Abcam, ab81887 or ab36001), anti-TUBB3 (1:200, Abcam, ab18207), anti-Ki67 (1:200, Fisher Scientific, 550609), anti-RAMP1 (1:200, Abcam, ab156575), anti-TROP2 (1:200, R&D Systems, AF1122), anti-DCLK1 (1:200, Cell Signaling Technology,62257S), anti-DCLK1 (1:200, Abnova,H00009201-M03), anti-CCK2R (1:200, Abcam, ab77077), anti-SUCNR1 (1:200, Novus Biologicals,NBP1-00861SS), anti-HPGDS (1:200, Abbexa,abx129473), and anti-POU2F3 (1:200, Atlas Antibodies,HPA019652). After washing, sections were incubated with Alexa Fluor-conjugated secondary antibodies (Alexa Fluor 488, 594, or 647; Thermo Fisher Scientific) for 1 h at room temperature. Nuclei were counterstained with DAPI. Images were quantified using Fiji (ImageJ; NIH). H&E-stained sections were analyzed using QuPath (v0.5.1) to delineate lesion areas, and dysplasia was graded by a gastrointestinal pathologist blinded to group allocation according to standard criteria.

**RNAscope in situ hybridization and bacterial FISH**

RNAscope in situ hybridization was performed using the RNAscope Multiplex Fluorescent Reagent Kit v2 (Advanced Cell Diagnostics) according to the manufacturer’s instructions. Paraffin-embedded gastric tissue sections were baked in a dry oven for 1 h at 60 °C followed by deparaffinization. Sections were then treated with hydrogen peroxide to block endogenous peroxidase activity. After hydrophobic barriers were drawn around the tissue sections, protease reagent was applied for target retrieval.

RNAscope probe targeting *P. melaninogenica*-16S rRNA (Advanced Cell Diagnostics, 841861) was applied to the sections and incubated in a humidified chamber at 40 °C. After probe hybridization, slides were washed twice in RNAscope wash buffer and sequential signal amplification was performed according to the manufacturer’s protocol. The signal was detected using Opal 570 (red channel). Sections were co-stained with anti-DCLK1 and anti-CGRP antibodies and DAPI to assess spatial proximity between bacteria, tuft cells, and sensory nerve endings.

**Lineage tracing**

For lineage tracing of CCK2R⁺ progenitors, *L2-IL1B; Cck2r-CreERT2; R26-tdTomato* mice received tamoxifen (100 mg kg⁻¹) to induce recombination, and GEJ tissues were collected at the indicated time points. The fraction of DCLK1⁺ tuft cells derived from CCK2R⁺ progenitors was quantified as the proportion of DCLK1⁺ cells expressing tdTomato on immunostained sections.

**AAV-mediated gene manipulation in the nodose ganglion**

For nodose ganglion (NG)-targeted viral delivery, mice were anesthetized with 2–4% isoflurane and placed on a heating pad. A ventral cervical incision was made, and the left NG was exposed under a stereomicroscope. A 33G Hamilton syringe was used to inject AAV vectors directly into the left NG. A total of 500 nl virus (1 × 10¹² vg ml⁻¹) or vehicle (PBS) was delivered slowly to minimize reflux and tissue damage, and the needle was left in place for 1–2 min before withdrawal. The incision was closed in layers, and mice were allowed to recover for 2–3 weeks before subsequent procedures.

For deletion of Sucnr1 in TRPV1-lineage vagal sensory neurons, rAAV-U6-sgSucnr1 (ABM) was injected into the left NG of *L2-IL1B; Trpv1-Cre; LSL-Cas9* mice; control animals received matched control vectors (AAV-Control) of the same serotype. Deletion efficiency was confirmed by immunostaining of NG sections.

Because viral delivery was consistently targeted to the left NG, tissue sampling for downstream histological analyses was standardized to the anterior walls of the stomach and esophagus.

**Monosynaptic retrograde rabies tracing**

Monosynaptic retrograde tracing was performed using an EnvA-pseudotyped glycoprotein-deleted rabies virus (EnvA-ΔG-RV) system.

To restrict rabies infection to defined gastric epithelial populations, a Cre-dependent helper AAV encoding TVA and optimized rabies glycoprotein (oG) (rAAV-DIO-Ef1a-TVA-2A-oG, 1 × 10¹² vg ml⁻¹, 10 μl per mouse; Addgene) was delivered to the anterior GEJ wall of *Cck2r-CreERT2* mice.

Three days after helper AAV delivery, mice were subjected to tamoxifen gavaging. One week after tamoxifen treatment, EnvA-ΔG-dsRed rabies virus (10⁸ IFU ml−1, 10 μl per mouse; Biohippo) was injected into the same GEJ region. Mice were euthanized 7 days later. The left NG was dissected and processed for fluorescence imaging to identify retrogradely labeled neurons.

**In vivo calcium imaging**

In vivo calcium imaging of tuft cells was performed in *L2-IL1B; Dclk1-CreERT2; R26-GCaMP6s* mice following tamoxifen induction. Mice were anesthetized with isoflurane and placed supine on a heated stage. The abdominal cavity was opened, and the GEJ region was exteriorized and stabilized for real-time microscopy. GCaMP6s signals were recorded with 488-nm excitation. Luminal succinate (10 mM in PBS) or vehicle was applied to the GEJ mucosa, and time-lapse images were acquired at a temporal resolution of 3 s. Responses were quantified as ΔF/F₀ relative to the pre-stimulus baseline in DCLK1-lineage (tdTomato⁺ or GCaMP6s⁺) tuft cells. Where indicated, imaging was compared between *P. melaninogenica*-colonized and control mice.

In vivo calcium imaging of gastric epithelial cells during vagal sensory neuron activation was performed using *L2-IL1B; Cck2r-CreERT2; R26-GCaMP6s; R26-tdTomato* mice. To enable optogenetic activation of vagal afferents, the left NG was injected with rAAV-Syn-ChrimsonR-tdTomato as described above, and experiments were performed 3 weeks later to allow robust expression. Mice were anesthetized with isoflurane and placed supine on a heated stage. The left NG was exposed by blunt dissection. For epithelial imaging, the abdominal cavity was opened, and the anterior GEJ wall was exteriorized and stabilized for real-time microscopy. Optogenetic stimulation was delivered to the exposed left NG using red light through a fiber-coupled light source (630–635 nm). Following 5 min of photostimulation of the left NG, time-lapse imaging of GCaMP6s fluorescence in Cck2r-lineage (tdTomato⁺) epithelial cells within the anterior GEJ was acquired in vivo. GCaMP6s signals were recorded with 488-nm excitation and appropriate emission collection, and tdTomato fluorescence was used to identify Cck2r-lineage cells.

**Ex vivo calcium imaging of nodose ganglion neurons**

Calcium imaging was performed on freshly isolated left NG from *Trpv1-Cre; R26-GCaMP6s* mice. NGs were immersed in Live Cell Imaging Solution (Thermo Fisher) and maintained at 37 °C during imaging. Time-lapse imaging was performed using a Leica TCS SP8 upright confocal microscope in XYZT acquisition mode at a temporal resolution of 3 s and a spatial resolution of 512 × 512 pixels. GCaMP6s fluorescence was excited at 488 nm and emission collected at 500–550 nm. After establishing a stable baseline, stimuli were applied by bath addition as indicated, including sodium succinate (10 mM), capsaicin (1 μM, Sigma-Aldrich), *P. melaninogenica* or *S. anginosus* conditioned medium (1:10 dilution) or their <3 kDa ultrafiltration fractions, and 15-d-PGJ2 (1 μM). For sensitization experiments, ganglia were pretreated with 15-d-PGJ2 or vehicle for 30 min before succinate stimulation. Regions of interest corresponding to individual TRPV1⁺ neurons were selected, and responses were quantified as ΔF/F₀ relative to the pre-stimulus baseline. Neurons were defined as responsive when ΔF/F₀ exceeded baseline fluctuations, and the fraction of responsive neurons and response amplitude were quantified for each condition.

**Nodose ganglion culture, neurite outgrowth, and CGRP release**

Left NGs were freshly isolated from adult mice (8–12 weeks old) and embedded in growth factor-reduced Matrigel (Corning) as 20-μl droplets in prewarmed 24-well plates. After polymerization for 20 min at 37 °C, wells were overlaid with culture medium. Recombinant mouse NGF (50 ng ml⁻¹), 15-d-PGJ2 (1 μM), sodium succinate (10 mM), bacterial conditioned medium or fractions, or vehicle were added as indicated. NGs were cultured for 3–5 days, then fixed and stained for TUBB3 to visualize neuronal processes. Axonal outgrowth was quantified using an outgrowth index calculated as the ratio of the axon-positive area fraction in the distal outgrowth region to that in the soma-adjacent region. For CGRP release assays, NG culture supernatants were collected at the indicated time points after stimulation, and CGRP levels were measured using a Mouse CGRP ELISA Kit (MyBioSource, MBS267121) according to the manufacturer's instructions.

**Flow cytometry and cell sorting**

For epithelial cell sorting, gastric epithelial cells were isolated from *Dclk1-CreERT2; R26-tdTomato* or *Cck2r-CreERT2; R26-ZsGreen* or *Dclk1-CreERT2; LSL-Cas9-EGFP* mice. Dead cells were excluded using DAPI, and epithelial lineage cells were identified as EpCAM⁺ (APC anti-mouse CD326/EpCAM, BioLegend, 118214) cells.All cell sorting experiments were performed using a BD Influx cell sorter.

**Organoid culture and coculture assays**

GEJ epithelial organoids were established from CCK2R⁺ progenitors isolated from *L2-IL1B; Cck2r-CreERT2; R26-tdTomato* mice. GEJ tissues were minced and digested in digestion medium (HBSS supplemented with 1% HEPES, 1 mg ml⁻¹ Collagenase P, 1 mg ml⁻¹ Dispase II, 10 μg ml⁻¹ DNase I, and 1% BSA) at 37 °C for 20 min with gentle agitation, filtered through a 40-μm strainer, and subjected to red blood cell lysis (BioLegend). Epithelial cells were stained with TruStain FcX (BioLegend) and APC anti-mouse CD326/EpCAM (BioLegend, 118214), and EpCAM⁺ tdTomato⁺ cells were sorted on a BD Influx cell sorter after exclusion of dead cells (DAPI).

A total of 2,000 sorted single epithelial cells were mixed with 20 μl growth factor-reduced Matrigel (Corning) per well and plated in 24-well plates. After Matrigel polymerization at 37 °C, organoid cultures were maintained in basal medium consisting of 50% conditioned medium containing Wnt3A, R-spondin-1, and Noggin (WRN, ATCC) and 50% Advanced DMEM/F12 (Gibco) supplemented with 20% FBS (Sigma-Aldrich), B27 (1×, Gibco), N2 (1×, Gibco), and 1% penicillin–streptomycin (Thermo Scientific). Medium was additionally supplemented with EGF (50 ng ml⁻¹, Gibco), FGF10 (100 ng ml⁻¹, PeproTech), Y-27632 (10 μM, Sigma-Aldrich), GlutaMAX (2 mM, Gibco), HEPES (10 mM), and SB431542 (10 μM, Tocris). Medium was replaced every other day. Organoids were passaged every 6 days at a 1:3 ratio by dissolving Matrigel with TrypLE (Gibco) followed by mechanical dissociation.

For NG–organoid cocultures, organoids were embedded in Matrigel domes and freshly isolated NGs were positioned adjacent to the dome (approximately 0.5–1 mm from the dome edge) and embedded in an additional Matrigel droplet. Cocultures were maintained in organoid growth medium, and organoid diameter and neuron–organoid contacts were quantified on day 5 unless otherwise indicated. Where indicated, DCLK1⁺ tuft cells isolated as described below were added to the coculture, and organoids were derived from *Ramp1⁺/⁺* or CCK2R-specific Ramp1-deficient (*Cck2r-CreERT2; Ramp1^flox/flox^*) mice. For chemogenetic stimulation of cocultured neurons, CNO (1 μM) was added to cultures of organoids with NGs from *Trpv1-Cre; R26-hM3Dq* mice.

**Tuft cell isolation, stimulation, and prostaglandin measurement**

DCLK1⁺ tuft cells were isolated from GEJ lesions of *P. melaninogenica*-colonized or control mice by FACS of EpCAM⁺ cells followed by FACS from *Dclk1-CreERT2; R26-tdTomato* reporter mice after tamoxifen induction. Sorted tuft cells were cultured for 48 h in organoid basal medium, and culture supernatants were collected. 15-d-PGJ2 and PGE2 concentrations in supernatants and in GEJ tissue lysates were quantified by ELISA (MyBioSource, MBS739421 and AFG Bioscience, EK248061) according to the manufacturer's instructions, normalized to cell number or tissue weight.

**CRISPR-mediated gene deletion in epithelial cells**

*Dclk1-CreERT2; LSL-Cas9-EGFP* mice received in vivo injections of an AAV vector carrying a Sucnr1-targeting sgRNA (sgSucnr1, ABM) or a control sgRNA (sgCtrl, ABM). Three weeks after AAV administration, mice were treated with tamoxifen to induce Cre-dependent Cas9-EGFP expression in Dclk1-lineage cells. One week after Cas9 induction, EGFP-positive epithelial cells were isolated by fluorescence-activated cell sorting (FACS) and co-cultured with tdTomato-labeled sensory neurons isolated from *Trpv1-Cre; R26-tdTomato* mice. Co-cultures were stimulated with succinate to assess the contribution of Sucnr1 in Dclk1-lineage tuft cells to succinate-induced neuronal responses.

**Statistical analysis**

Statistical analyses were performed using GraphPad Prism (v.10) and Python. Data are presented as mean ± s.e.m. unless otherwise indicated. For quantification of cell numbers and nerve densities, measurements from multiple fields or sections were averaged for each mouse, and individual mice were treated as biological replicates. Unless otherwise specified, comparisons between two independent groups were performed using unpaired two-tailed Student’s t-tests. Comparisons among multiple groups were performed using one-way or two-way ANOVA followed by the post hoc tests specified in the corresponding figure legends. Categorical variables were analyzed using Fisher’s exact test or the chi-square test, as appropriate. For microbiome analyses, paired comparisons of alpha diversity were performed using two-tailed Wilcoxon signed-rank tests, differences in beta diversity were assessed using PERMANOVA, and differentially abundant taxa were identified using LEfSe. For metabolomic analyses, P values were adjusted for multiple testing using the Benjamini–Hochberg method. For scRNA-seq analyses, differential gene expressions were assessed using Wilcoxon rank-sum tests with Bonferroni correction. Additional analyses, including Friedman tests, Spearman correlations, and linear mixed-effects models, are described in the relevant Methods sections and figure legends. A *P* value < 0.05 was considered statistically significant; where multiple-testing correction was applied, significance was determined using adjusted P values. Statistical significance is indicated as follows: ns, not significant; **P* < 0.05; ***P* < 0.01; ****P* < 0.001; and *****P* < 0.0001. Sample sizes, experimental units, and statistical tests are specified in the corresponding figure legends.

**Supplementary Figures**

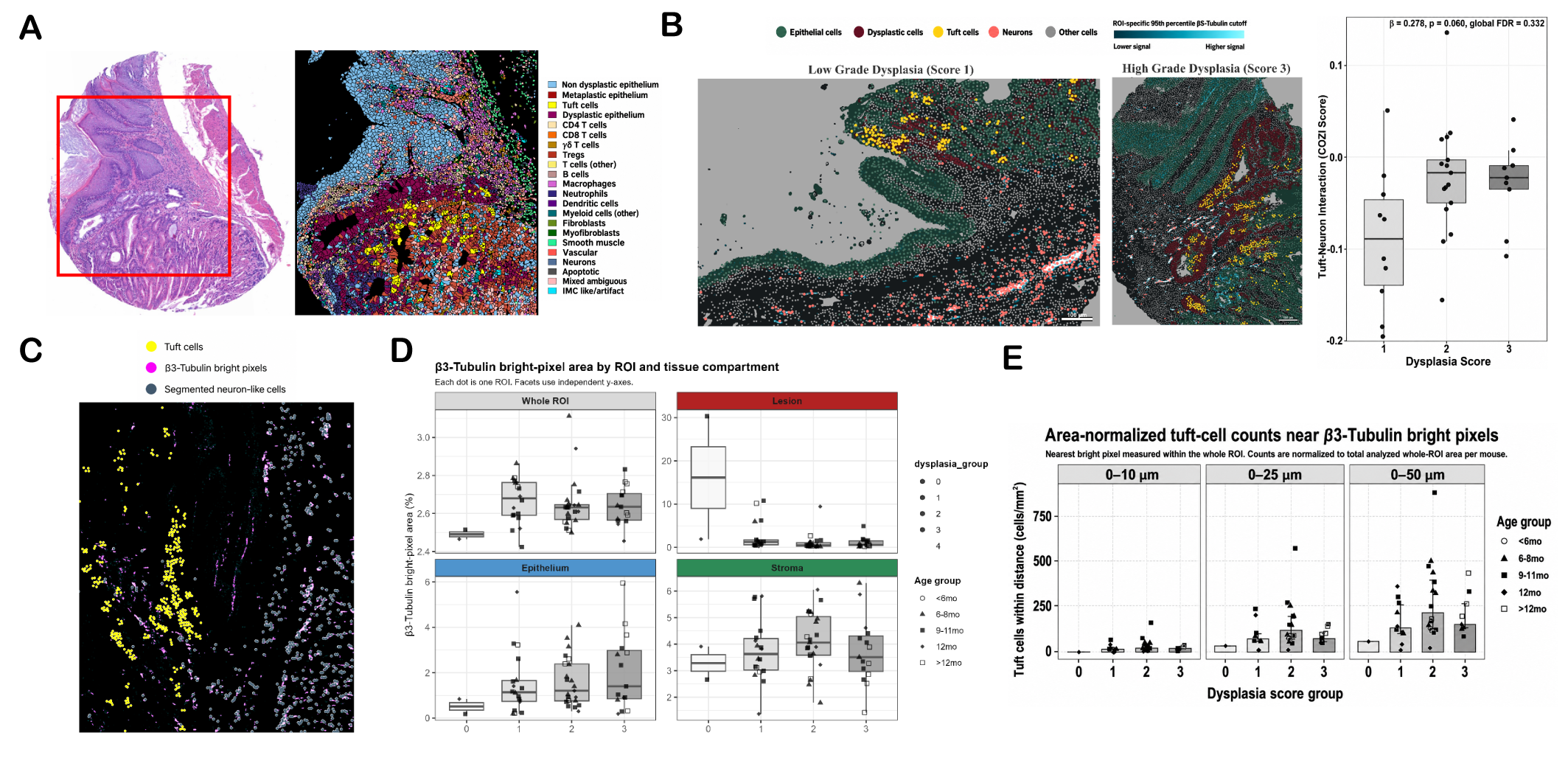

**Fig. S1. Spatial analysis of tuft cell–neuron proximity by imaging mass cytometry (supports Fig. 1).**

(**A**) Representative H&E-stained GEJ section and corresponding IMC cell-type map showing the full phenotypic annotation of segmented cells. (**B**) Directional neighbor-preference analysis of tuft cell–neuron spatial interactions. Representative spatial maps of low-grade (score 1) and high-grade (score 3) dysplasia regions of interest (ROIs) are shown together with tuft cell→neuron COZI interaction scores across dysplasia score groups. The interaction increased with dysplasia severity but did not remain significant after multiple-testing correction (linear mixed-effects coefficient β = 0.278, nominal *P* = 0.060, global FDR = 0.332).(**C**) Representative continuous β3-tubulin image-field analysis showing DCLK1+ tuft cells (yellow), β3-tubulin–bright pixels (magenta), and segmented neuron-like cells (blue).(**D**) β3-tubulin–bright-pixel area fraction in the whole ROI and within lesion, epithelial, and stromal compartments across dysplasia score groups. Each point represents one ROI. (**E**) Area-normalized tuft-cell counts within 0–10, 0–25, and 0–50 μm of β3-tubulin–bright structures across dysplasia score groups. Counts were normalized to total analyzed whole-ROI area per mouse.

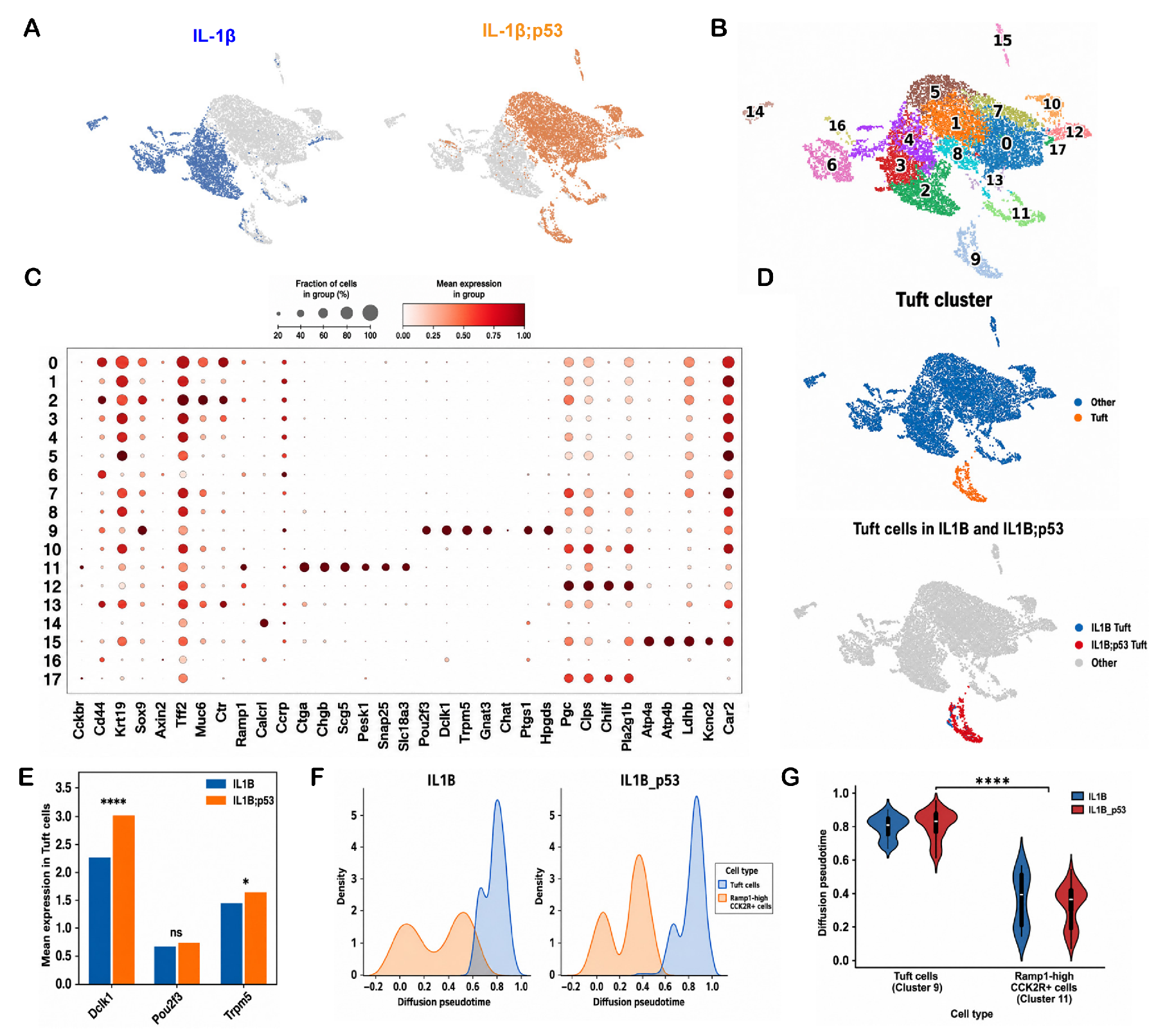

**Fig. S2. Single-cell RNA sequencing of mouse GEJ epithelium (supports Fig. 2).**

(**A**) UMAP embeddings of GEJ epithelial cells from IL-1β and IL-1β; p53 mice (GSE246280). (**B**) UMAP with cluster identities (clusters 0–17). (**C**) Dot plot of lineage-marker gene expression across all clusters. (**D**) UMAP highlighting the tuft-cell cluster (top) and tuft cells by genotype (bottom). (**E**) Mean expression of Dclk1, Pou2f3, and Trpm5 in tuft cells by genotype. (**F**) Density distributions of diffusion pseudotime for tuft cells and Ramp1-high CCK2R+ cells within each genotype. (**G**) Violin plots of diffusion pseudotime for tuft cells (cluster 9) and Ramp1-high CCK2R+ cells (cluster 11) by genotype. For all panels, values indicate mean ± SEM. **P* < 0.05, ***P* < 0.01, ****P* < 0.001, *****P* < 0.0001; ns: not significant.

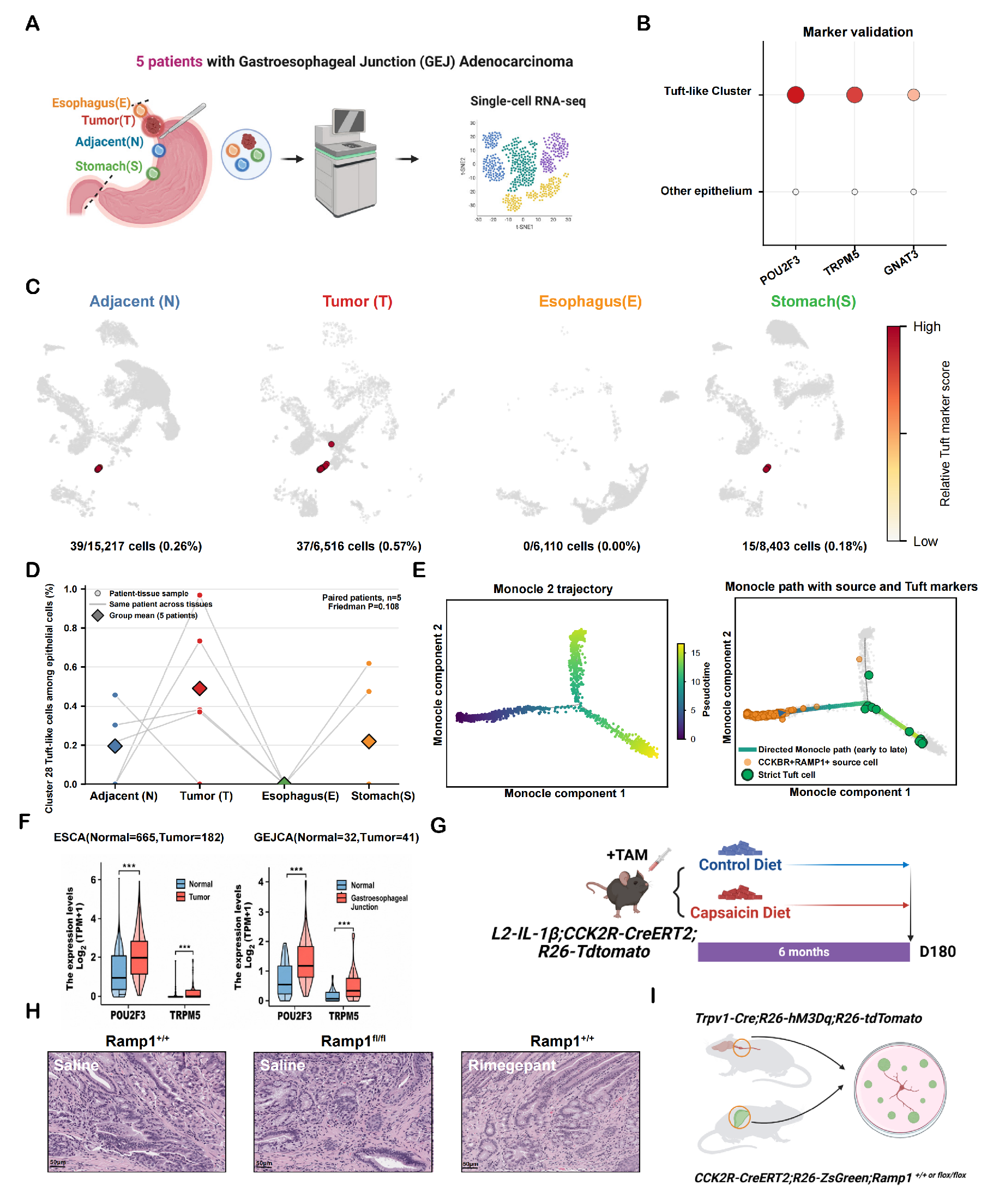

**Fig. S3. Single-cell analysis of human GEJ adenocarcinoma and validation cohorts (supports Fig. 2).**

(**A**) Schematic: single-cell RNA sequencing of four anatomical sites (esophagus, tumor, adjacent, stomach) from 5 patients with GEJ adenocarcinoma. (**B**) Marker validation: POU2F3, TRPM5, and GNAT3 expression in the tuft-like cluster versus other epithelium. (**C**) UMAP maps of tuft-marker scores by anatomical site, with tuft-like cell counts and frequencies (adjacent 39/15,217 cells, 0.26%; tumor 37/6,516, 0.57%; esophagus 0/6,110, 0.00%; stomach 15/8,403, 0.18%). (**D**) Paired analysis of tuft-like cell frequencies across sites within the same patients (n = 5; Friedman test, *P* = 0.109). (**E**) Monocle 2 trajectory of epithelial cells colored by pseudotime (left) and the directed Monocle path from CCK2R+RAMP1+ source cells to strict tuft cells (right). (**F**) POU2F3 and TRPM5 expression in normal versus tumor tissues in TCGA ESCA (normal = 665, tumor = 182) and a GEJ adenocarcinoma cohort (normal = 32, tumor = 41). (**G**) Experimental scheme for chronic capsaicin feeding: *L2-IL1B; Cck2r-CreERT2; R26-tdTomato* mice on control or capsaicin diet for 6 months (supports Fig. 2H). (**H**) H&E-stained GEJ sections from *Ramp1^+/+^* + saline, *Ramp1^fl/fl^* + saline, and *Ramp1^+/+^* + rimegepant mice (supports Fig. **2, I** and **J**). (**I**) Schematic of the NG–organoid coculture system: NG from Trpv1-Cre; R26-hM3Dq; R26-tdTomato mice with CCK2R+ organoids from *Cck2r-CreERT2; R26-ZsGreen; Ramp1^+/+^* or *Ramp1^fl/fl^* mice (supports Fig. **2K**). Scale bars: 50 μm (**H**). For all panels, values indicate mean ± SEM. **P* < 0.05, ***P* < 0.01, ****P* < 0.001, *****P* < 0.0001; ns: not significant. Schematic in (**A, G, I**) created with BioRender.com.

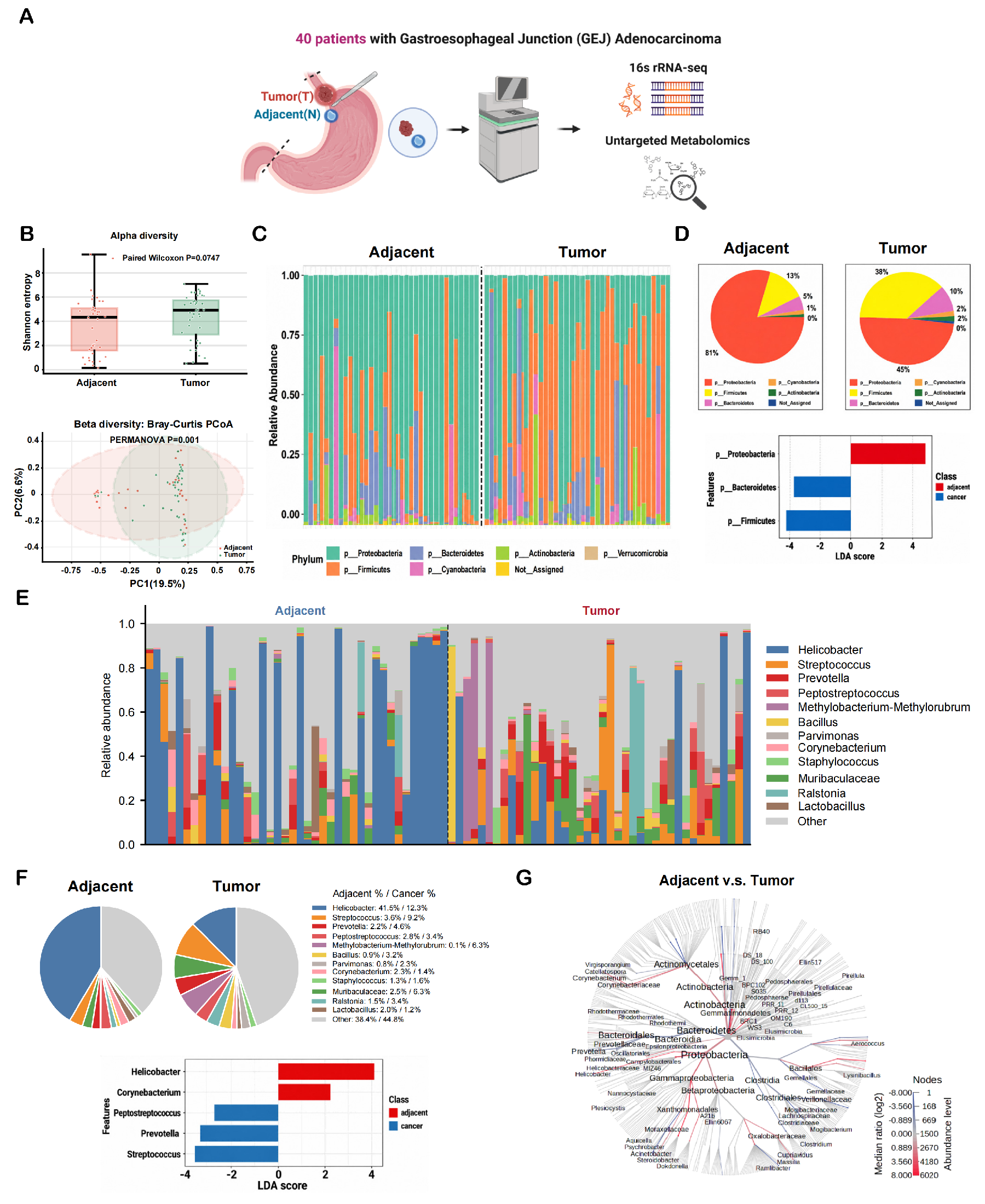

**Fig. S4. Microbiome profiling of the human GEJ adenocarcinoma cohort (supports Fig. 3).**

(**A**) Schematic: paired tumor and adjacent tissues from 40 patients with GEJ adenocarcinoma analyzed by 16S rRNA sequencing and untargeted metabolomics. (**B**) Alpha diversity (Shannon entropy; paired Wilcoxon test, *P* = 0.0747) and beta diversity (Bray–Curtis PCoA; PERMANOVA, *P* = 0.001) of adjacent versus tumor microbiota. (**C**) Phylum-level relative-abundance stacked bars for all paired samples. (**D**) Phylum-level composition (pie charts) and LEfSe LDA scores at phylum level. (**E**) Genus-level relative-abundance stacked bars. (**F**) Genus-level composition with adjacent/tumor percentages and LEfSe LDA scores (Helicobacter and Corynebacterium enriched in adjacent tissue; Peptostreptococcus, Prevotella, and Streptococcus enriched in tumor). (**G**) Taxonomic tree of adjacent- versus tumor-associated taxa (node color, median log2 ratio; node size, abundance level). Schematic in (**A**) created with BioRender.com.

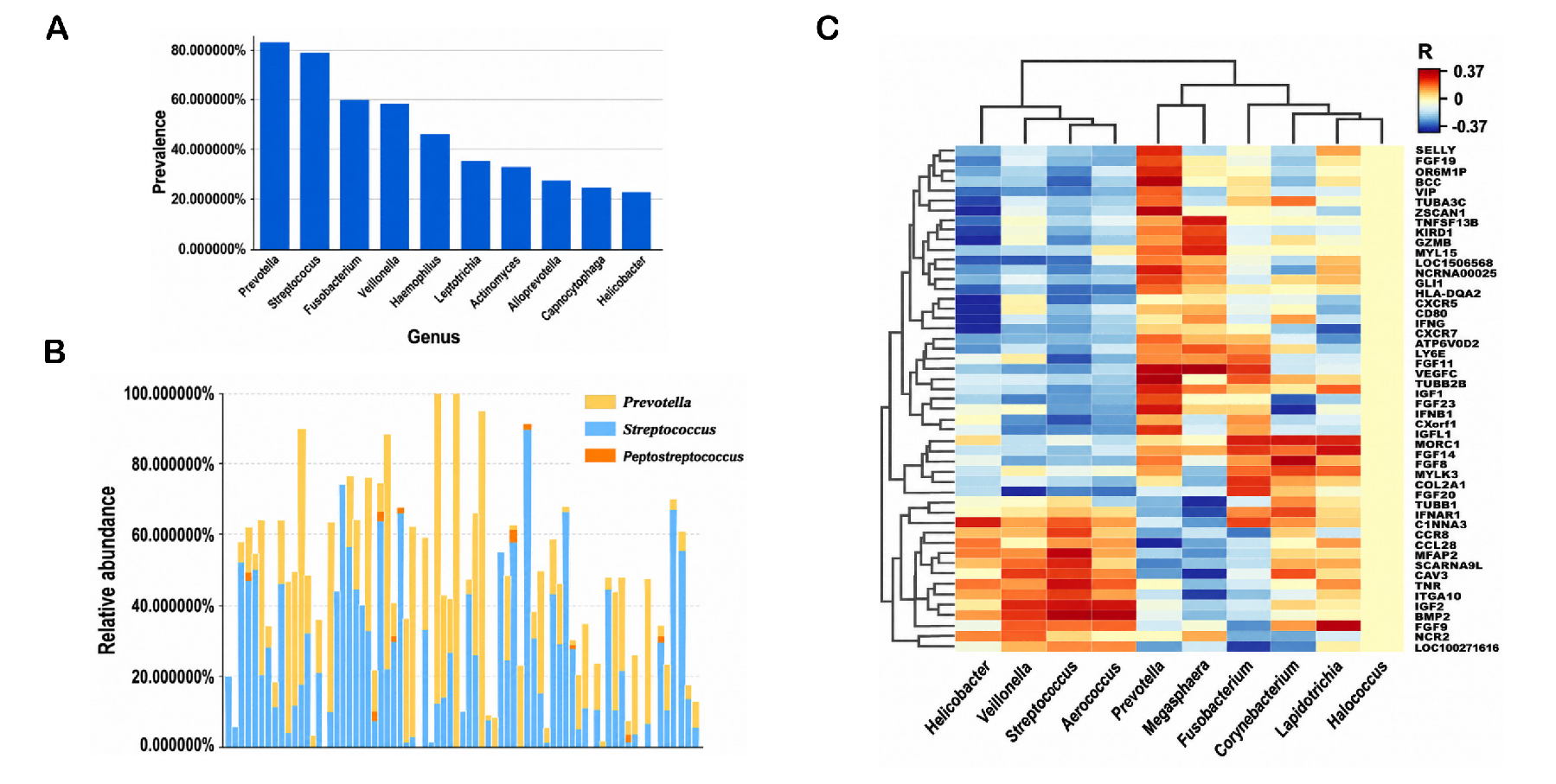

**Fig. S5. Validation of tumor-associated microbial taxa and neuronal gene correlations in TCMA (supports Fig. 3).**

(**A**) Prevalence of tumor-associated bacterial genera in The Cancer Microbiome Atlas (TCMA) cohort. (**B**) Relative abundance of *Prevotella*, *Streptococcus*, and *Peptostreptococcus* across individual TCMA samples. (**C**) Hierarchically clustered heatmap of correlations between genus abundance and tumor gene expression, highlighting positive associations of *Prevotella* with neuronal development and neurite-related genes, including TUBB2B, FGF11, and FGF14.

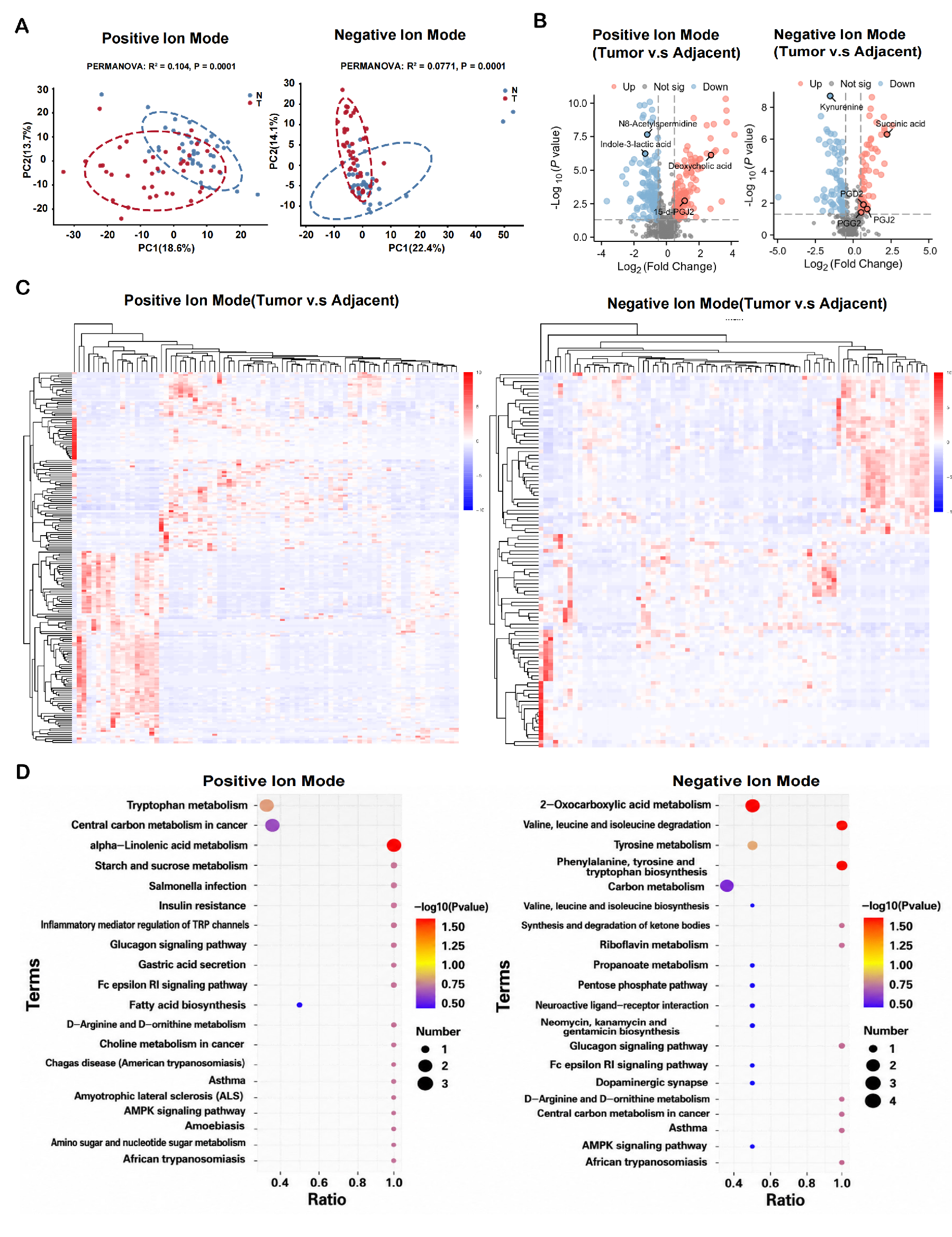

**Fig. S6. Untargeted metabolomics of the human GEJ adenocarcinoma cohort (supports Fig. 3).**

(**A**) PCA of metabolomic profiles in positive (PERMANOVA R² = 0.104, *P* = 0.0001) and negative (R² = 0.0771, *P* = 0.0001) ion modes, adjacent (N) versus tumor (T). (**B**) Volcano plots of differential metabolites (tumor versus adjacent) in both ion modes; labeled metabolites include deoxycholic acid, succinic acid, and 15-d-PGJ2 (up in tumor) and N8-acetylspermidine, indole-3-lactic acid, PGD2, PGG2 and PGJ2 (down or unchanged). (**C**) Hierarchically clustered heatmaps of metabolite abundances in both ion modes. (**D**) KEGG pathway enrichment of differential metabolites in both ion modes (bubble size, metabolite number; color, −log10 *P*).

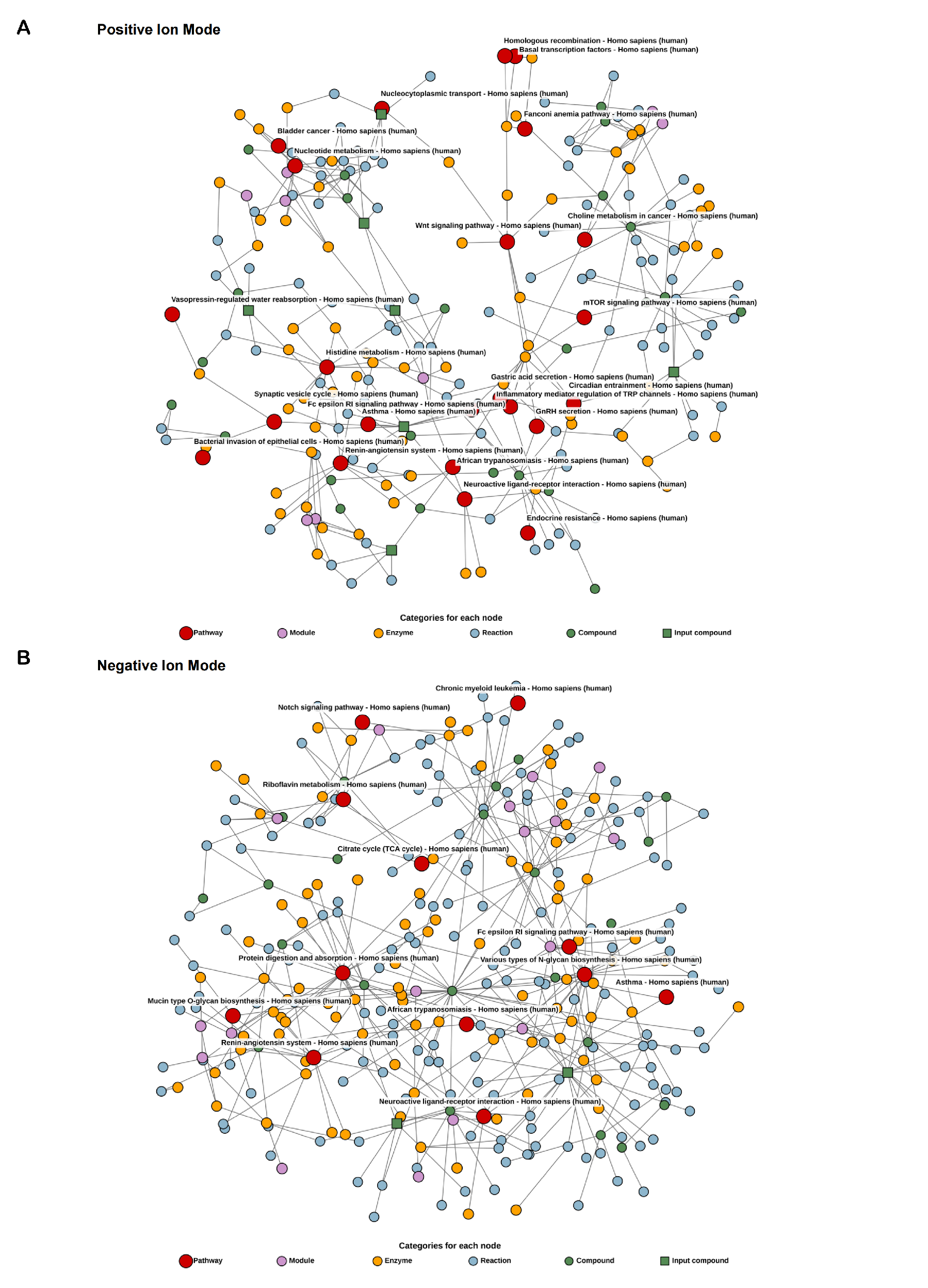

**Fig. S7. KEGG pathway–compound networks of differential metabolites (supports Fig. 3).**

(**A**) Positive ion mode and (**B**) negative ion mode network graphs linking differential metabolites (input compounds) to enzymes, reactions, modules, compounds, and pathways.

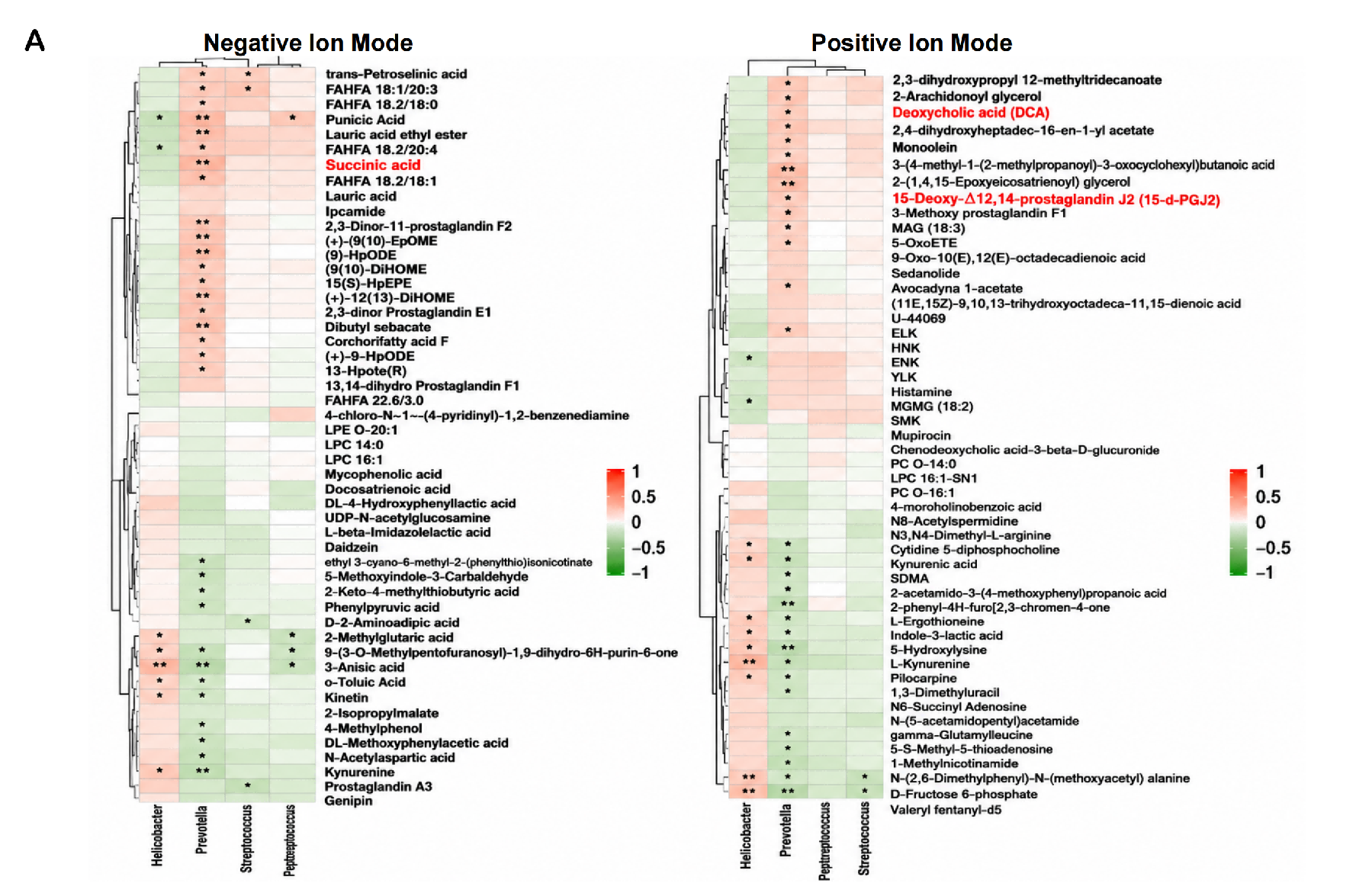

**Fig. S8. Correlation between tumor-associated genera and metabolites (supports Fig. 3).**

(**A**) Hierarchically clustered heatmaps of correlations between the four dominant genera (*Helicobacter, Prevotella, Streptococcus, Peptostreptococcus*) and metabolite abundances in negative (left) and positive (right) ion modes; succinic acid, deoxycholic acid (DCA), and 15-deoxy-Δ12,14-prostaglandin J2 (15-d-PGJ2) are highlighted in red. **P* < 0.05, ***P* < 0.01.

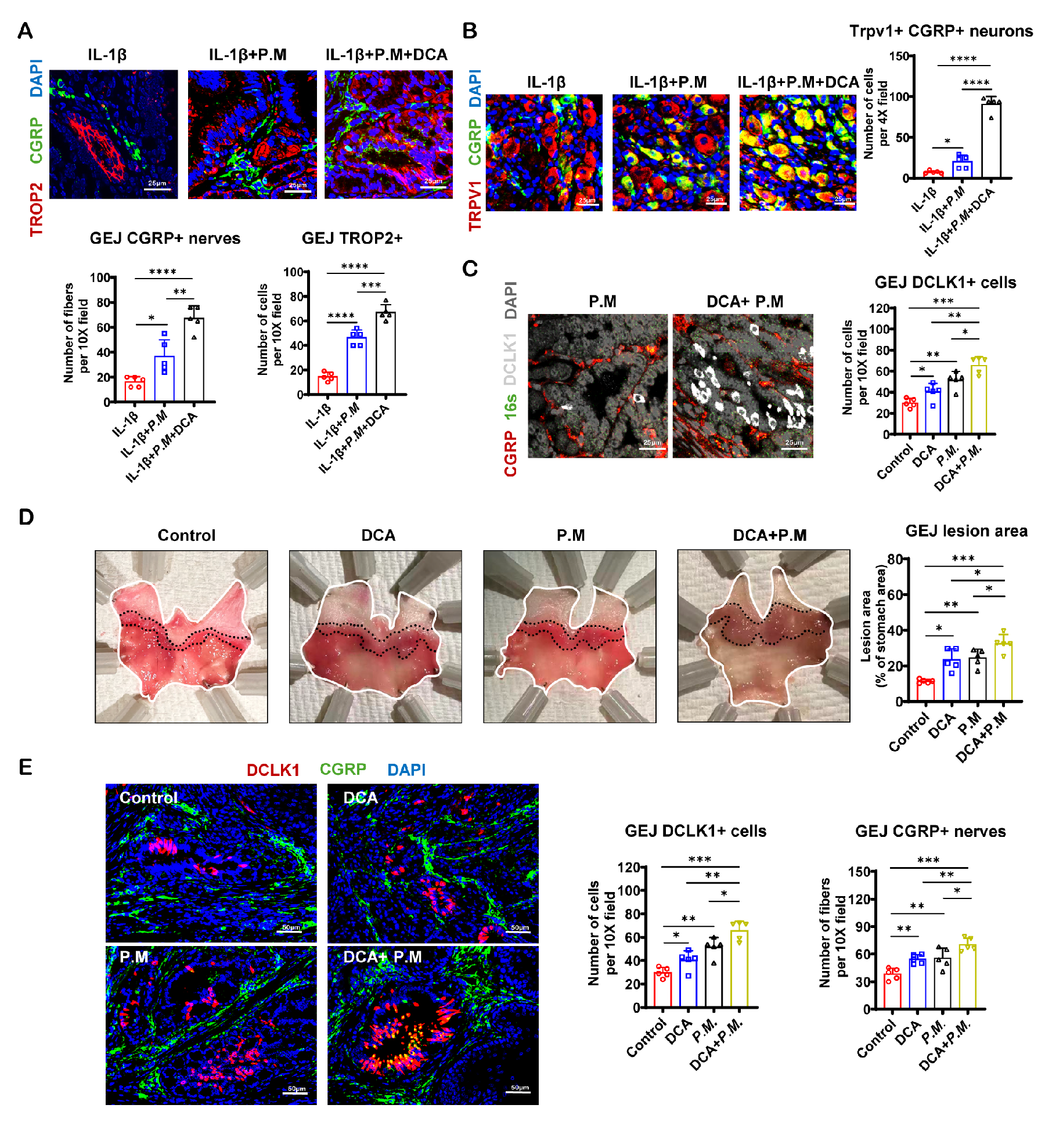

**Fig. S9. *P. melaninogenica* and DCA cooperatively remodel the GEJ neuroepithelial niche in vivo (supports Fig. 3).**

(**A**) IF of CGRP and TROP2 at the GEJ of *L2-IL1B*, *L2-IL1B* + *P. melaninogenica*, and *L2-IL1B* + *P. melaninogenica* + DCA mice, with quantification of CGRP+ nerves and TROP2+ cells. n = 5 mice per group. (**B**) IF of CGRP and TRPV1 in nodose ganglia of the same groups, with quantification of TRPV1+CGRP+ neurons per 4× field. n = 5 mice per group. (**C**) RNAscope for *P. melaninogenica* 16S rRNA combined with DCLK1 and CGRP IF in colonized mice with or without DCA, with quantification of GEJ DCLK1+ cells across control, DCA, *P. melaninogenica*, and DCA + *P. melaninogenica* groups. n = 5 mice per group. (**D**) Representative whole-mount stomachs of the four groups with quantification of GEJ lesion area. n = 5 mice per group. (**E**) IF of DCLK1 and CGRP in the four groups, with quantification of DCLK1+ cells and CGRP+ nerves. n = 5 mice per group. Scale bars: 50 μm (**A, C, E**), 20 μm (**B**). For all panels, values indicate mean ± SEM. **P* < 0.05, ***P* < 0.01, ****P* < 0.001, *****P* < 0.0001; ns: not significant.

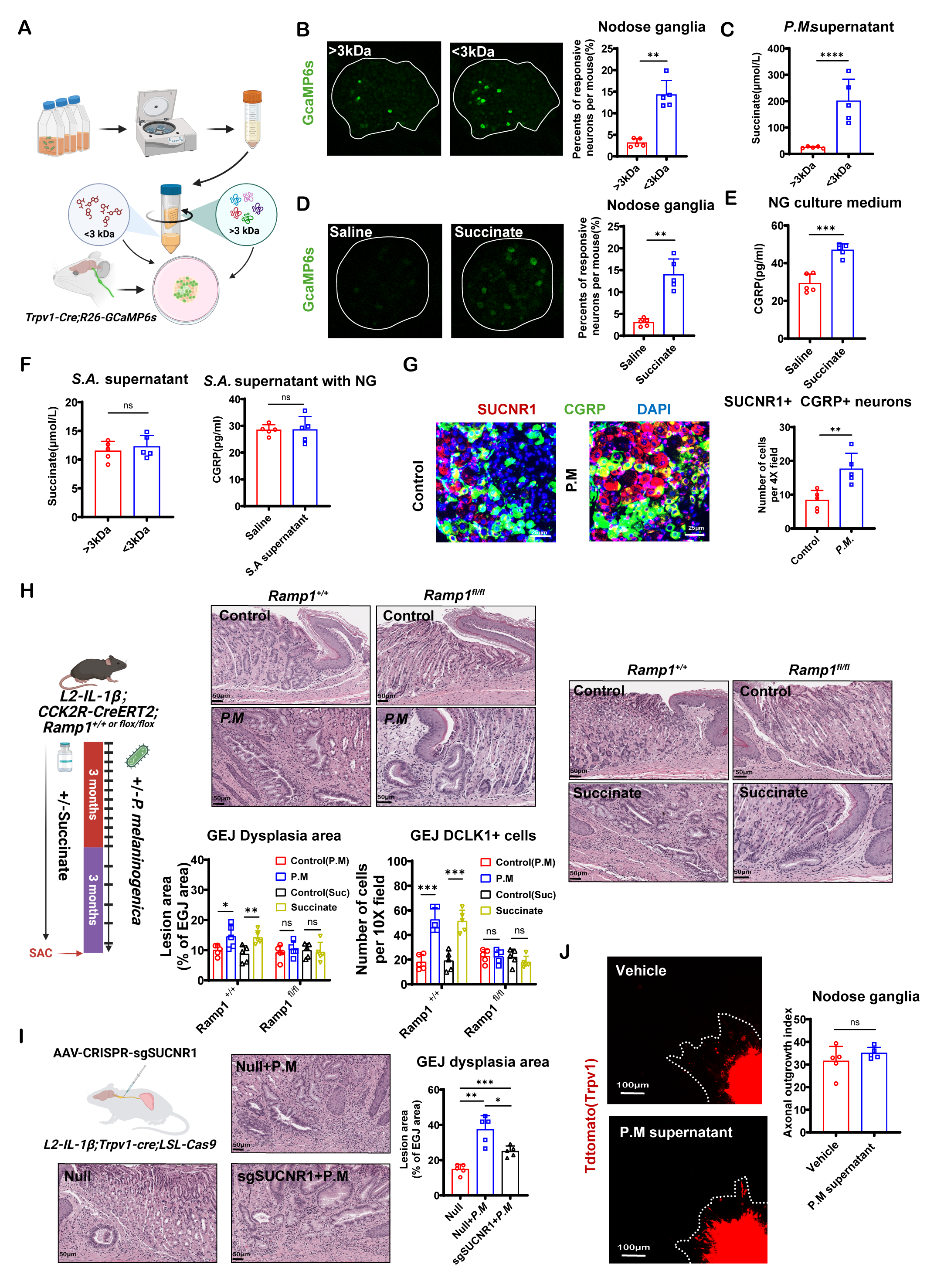

**Fig. S10. Succinate mediates *P. melaninogenica* activation of vagal sensory neurons (supports Fig. 4).**

(**A**) Schematic: *P. melaninogenica*–conditioned medium fractionated by 3-kDa cutoff ultrafiltration and applied to nodose ganglia from *Trpv1-Cre; R26-GCaMP6s* mice. (**B**) GCaMP6s responses of nodose ganglion neurons to >3-kDa versus <3-kDa fractions, with quantification of responsive neurons per mouse. n = 5 mice per group. (**C**) Succinate concentrations in >3-kDa versus <3-kDa fractions of *P. melaninogenica* supernatant. n = 5 mice per group. (**D**) GCaMP6s responses to saline versus succinate, with quantification. n = 5 mice per group. (**E**) CGRP release into nodose ganglion culture medium after saline versus succinate stimulation. n = 5 mice per group. (**F**) Specificity control: succinate content of *Streptococcus anginosus* (S.A.) supernatant fractions (left) and CGRP release from nodose ganglia exposed to S.A. supernatant (right). n = 5 mice per group. (**G**) IF of SUCNR1 and CGRP in nodose ganglia of control versus colonized mice, with quantification of SUCNR1+CGRP+ neurons per 4× field. n = 5 mice per group. (**H**) Experimental scheme: *L2-IL1B; Cck2r-CreERT2; Ramp1^+/+^* or *Ramp1^fl/fl^* mice treated with succinate (100 mM in drinking water) and/or *P. melaninogenica* colonization over 6 months, wtih H&E staining and quantification of GEJ dysplasia area and DCLK1+ cells.n = 5 mice per group. (**I**) AAV-CRISPR–mediated Sucnr1 deletion in TRPV1-lineage nodose ganglion neurons (*L2-IL1B; Trpv1-Cre; LSL-Cas9*): H&E sections of Null, Null + *P. melaninogenica*, and sgSUCNR1 + *P. melaninogenica* groups, with quantification of GEJ dysplasia area.n = 5 mice per group. (**J**) Neurite outgrowth from *Trpv1-Cre; R26-tdTomato* nodose ganglia exposed to vehicle or *P. melaninogenica* supernatant; axonal outgrowth index. n = 5 mice per group. Scale bars: 50 μm (**G–I**), 100 μm (**J**). For all panels, values indicate mean ± SEM. **P* < 0.05, ***P* < 0.01, ****P* < 0.001, *****P* < 0.0001; ns: not significant. Schematic in (**A, H, I**) created with BioRender.com.

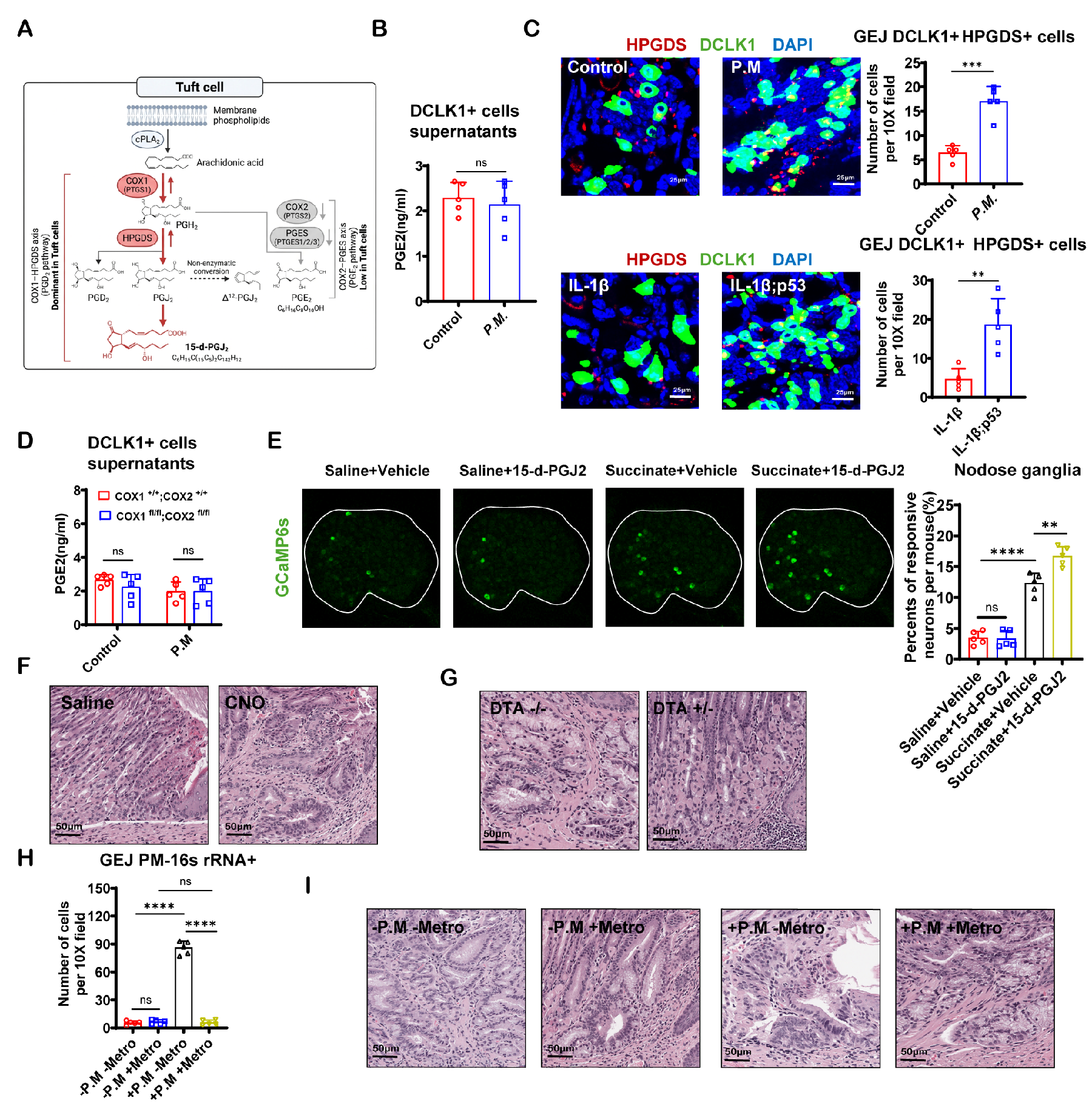

**Fig. S11. The HPGDS/15-d-PGJ2 axis in tuft cells (supports Fig. 5).**

(**A**) Schematic of prostaglandin synthesis in tuft cells: the COX1 (PTGS1)–HPGDS axis (PGD2 pathway, dominant in tuft cells) generates PGD2, PGJ2, and 15-d-PGJ2, whereas the COX2 (PTGS2)–PTGES axis (PGE2 pathway) is minimally expressed in tuft cells. (**B**) PGE2 concentrations measured by ELISA in culture supernatants of DCLK1⁺ tuft cells sorted from control or *P. melaninogenica*–colonized mice. n = 5 mice per group. (**C**) IF of HPGDS and DCLK1 at the GEJ of control versus colonized mice (top) and IL-1β versus IL-1β; p53 mice (bottom), with quantification of DCLK1+HPGDS+ cells. n = 5 mice per group. (**D**) PGE2 concentrations measured by ELISA in culture supernatants of DCLK1⁺ tuft cells of Cox1/Cox2-intact versus tuft-specific Cox1/Cox2-deficient mice, with or without colonization (all ns). n = 5 mice per group. (**E**) GCaMP6s responses of nodose ganglion neurons to saline or succinate, with vehicle or 15-d-PGJ2 pretreatment, with quantification of responsive neurons per mouse. n = 5 mice per group. (**F**) H&E-stained GEJ sections from saline- versus CNO-treated *L2-IL1B; Dclk1-CreERT2; R26-hM3Dq* mice (supports Fig. **5, F** and **G**). (**G**) H&E-stained GEJ sections from *L2-IL1B; Pou2f3-CreERT2; R26-DTA DTA*^−/−^ versus *DTA^+/−^* mice (supports **Fig. 5H**). (**H**) Quantification of GEJ *P.M*-16S rRNA+ cells across the four metronidazole/colonization groups (supports **Fig. 5I**). n = 5 mice per group. (**I**) H&E-stained GEJ sections of the four groups (supports **Fig. 5I**). Scale bars: 25 μm (**C**), 50 μm (**F, G, I**). For all panels, values indicate mean ± SEM. **P* < 0.05, ***P* < 0.01, ****P* < 0.001, *****P* < 0.0001; ns: not significant. Schematic in (**A**) created with BioRender.com.
